# LncRNA MALAT1/microRNA-30b axis regulate macrophage polarization and function

**DOI:** 10.1101/2023.02.01.526668

**Authors:** Imran Ahmad, Raza Ali Naqvi, Araceli Valverde, Afsar R. Naqvi

## Abstract

**Introduction:** Macrophages (Mφ) can polarize towards the proinflammatory M1 or proresolving M2 phenotype to control diverse biological processes such as inflammation, and tissue regeneration. Noncoding RNAs play critical roles in numerous biological pathways; however, their functional interaction in the regulation of Mφ polarization and immune responses remain unclear.

**Objectives:** To examine relationship between lncRNA (MALAT1) and microRNA (miR-30b) in shaping macrophage polarization and immune functions.

**Methods:** Expression of MALAT1 and miR-30b was examined in differentiating M1/M2 Mφ, human and murine inflamed gingival biopsies by RT-qPCR. MALAT1 and miR-30b direct interaction was examined by dual luciferase assays. Impact of MALAT1 knockdown and miR-30b overexpression was examined on macrophage polarization markers, bacterial phagocytosis, antigen uptake/processing and cytokine profiles.

**Results:** MALAT1 expression displays a time-dependent induction during Mφ differentiation and, upon challenge with TLR4 agonist (*E. coli* LPS). Knockdown of MALAT1 enhanced the expression of M2Mφ markers without affecting the M1Mφ markers, suggesting that MALAT1 favors the M1 phenotype by suppressing M2 polarization. MALAT1 knockdown Mφ exhibit reduced antigen uptake and processing, bacterial phagocytosis, and bactericidal activity, strongly supporting its critical role in regulating innate immune functions. Consistent with this, MALAT1 knockdown showed impaired cytokine secretion upon challenge with LPS. Importantly, MALAT1 exhibit an antagonistic expression pattern with all five members of the miR-30 family during M2Mφ differentiation. Dual-luciferase assays validated a novel sequence on MALAT1 that interacts with miR-30b, a microRNA that promotes the M2 phenotype. Phagocytosis and antigen processing assays unequivocally demonstrated that MALAT1 and miR-30b are functionally antagonistic. In human subjects with periodontal disease and murine model of ligature-induced periodontitis, we observed higher levels of MALAT1, and downregulation of miR-30b that correlates with higher M1Mφ markers expression in gingival tissues suggesting a pro-inflammatory function of MALAT1.

**Conclusion:** MALAT1/miR-30b antagonistic interaction shapes Mφ polarization *in vitro* and in inflamed gingival biopsies.

## Introduction

The ability of a healthy immune system to clear a plethora of antigens relies on the enormous plasticity displayed by the relevant cell types. Monocytes, macrophages (Mφ), and dendritic cells (DCs) are members of the mononuclear phagocyte system (MPS) that constantly patrol peripheral tissues. These cells are the imperative players in innate immunity, which form the first lines of defense and are responsible for maintaining tissue integrity and homeostasis via active recruitment to sites of injury and infection [1]. Mφ recognize pathogens by diverse Toll-like receptors (TLRs), phagocytize them, secrete cytokines that eventually activate the adaptive arm of the immune system, and perform critical roles in wound repair. The coordinated combination of local stimuli facilitates the activation of differential switches inside Mφ and induces them to acquire specialized functional phenotypes, viz. inflammatory (M1) and reparative (M2) subtypes, depending upon the phase of infection [1–3]. Mφ polarization is a tightly controlled process, and numerous endogenous and exogenous molecules can skew their phenotype. Dysregulation of Mφ polarization has been implicated in autoimmune diseases, cancers, fibrosis, viral infections, etc. [3–8]. However, the mechanistic details of this process and the molecular events regulating Mφ polarization are not fully understood. Therefore, deciphering the role of endogenous regulatory molecules that control the differentiation and function of these multifunctional immune cells can uncover novel regulatory mechanisms.

One intriguing finding of human genome annotation was the abundance of transcripts with little or no protein-coding potential, later termed long (>200 nts) noncoding RNAs (lncRNAs) [9, 10]. As of now, more than 50,000 different lncRNAs have been identified in humans, and the list continues to expand [11]. However, a large repertoire of lncRNAs remains uncharacterized. Interestingly, lncRNAs are modular molecules that can physically interact with DNA, RNA (mRNA/miRNA) or protein to regulate transcriptional, posttranscriptional and translational output of the genome [12–14]. Aberrations in lncRNA expression have been reported to be associated with disease manifestation in cancers, neurodegenerative disorders, autoimmunity, etc. [15].

Macrophage polarization is a dynamic process that is regulated by changes in gene expression, transcription factors, epigenetic modifications, signaling pathways, and posttranslational modifications [16–18]. Recent studies have uncovered the differential expression of lncRNAs during macrophage differentiation and macrophage polarization in various diseases [19–22]. Currently, few lncRNAs have been demonstrated to regulate macrophage polarization. For instance, lncRNAs AFAP-AS1 and PVT1 promote M1 polarization, while DLX6-AS1 and LINC00467 promote M2 polarization [23–25]. However, lncRNA-mediated regulation of Mφ phenotype and innate immune functions and their interaction with other noncoding RNAs during these biological processes remain poorly studied.

A recent publication from our laboratory methodically showed the differential lncRNA expression profile in monocyte-to-macrophage differentiation, and we observed higher expression of MALAT1 [26]. In this article, we examined the role of MALAT1 in macrophage polarization and the underlying mechanisms, which have not been fully explored. Furthermore, we investigated MALAT1-mediated regulation of macrophage polarization through microRNA sponging. Our results show that lncRNA MALAT1 and miR-30b crosstalk was instrumental in skewing M1/M2 polarization and regulating key innate immune functions including phagocytosis and antigen processing. We examined MALAT1 and miR-30b profiles and their relationship with macrophage polarization markers in gingival (gum) biopsies collected from periodontally diseased human subjects and mice model of ligature-induced periodontitis.

## Materials and Methods

### Primary human monocyte isolation and differentiation

The freshly prepared buffy coats collected from healthy donors were used to purify Peripheral blood mononuclear cells (PBMCs) (Sylvan N. Goldman Oklahoma Blood Institute, Oklahoma City, OK) using Ficoll Paque (GE Healthcare, Piscataway, NJ)-based density centrifugation method, as described earlier [27, 28]. The purified PBMCs were used to isolate CD14+ monocytes, which were differentiated into M1 Mφ, M2 Mφ or DCs in the presence of recombinant human (rh) GM-CSF (1000 U/ml), M-CSF (50 ng/ml) or GM-CSF+IL-4 (50 ng/ml; PeproTech, Rocky Hill, NJ). Cells were harvested at various time points for flow cytometric analysis or RNA isolation.

### Transient GapmeR and miRNA transfection

MALAT1 and control GapmeRs were purchased from Qiagen (Germantown, MD). Transient transfection of GapmeRs was performed using Lipofectamine 2000 (Life Technologies) according to the manufacturer’s instructions. Cells were transfected with lncRNA MALAT1 or control GapmeR at a final concentration of 50 nM. Red siGLO oligos (Thermo Scientific, Waltham, MA) were used to determine transfection efficiency.

### Cell viability assay

Cell viability assays were performed using the CellTiter 96 AQueous Cell Proliferation Assay Kit (Promega, Madison, WI), according to the manufacturer’s protocol. Briefly, CD14+ monocytes were seeded and differentiated in 96-well plates and transfected with MALAT1 or control GapmeR, and assays were performed at 24 h and 48 h post transfection

### Total RNA isolation, cDNA synthesis and qPCR

Total RNA was isolated from monocytes, M1/M2 Mφ and DCs using the miRNeasy mini-kit (Qiagen) according to the manufacturer’s instructions. 500 ng of total RNA was used to synthesize first-strand cDNA using the Reverse Transcription Kit (Qiagen). For lncRNA expression profiling, commercially available primers were purchased from Sigma, and transcript expression was quantified by real-time PCR using a StepOne plus thermocycler (Applied Biosystems, Carlsbad, CA). Expression levels were normalized with respect to β-actin, and the fold change was calculated using the delta-delta CT method. M1 or M2 Mφ were harvested at 36 h post-transfection with lncRNA or treatment with LPS. Total RNA was isolated, and the expression of MALAT1 lncRNA and gene markers of M1 (iNOS, STAT1, TNF-α, ARG2) and M2 (ARG1, STAT3, CCL2, IL-10) Mφ was analyzed by RT–qPCR. The data are presented as normalized fold change with SD.

### Flow cytometry

Cells were harvested and washed in ice-cold PBS supplemented with 1% (v/v) FBS and 0.08% sodium azide. Samples were analyzed using a FACScan or BD Cyan flow cytometer using CellQuest software (BD Biosciences, San Jose CA). Equal number of cells (10,000 or 20,000) were captured for all the experiments. Cellular debris was excluded based on size (forward scatter [FSC]) and granularity (side scatter [SSC]). Couplets were excluded based on SSC versus FSC and SSC versus pulse width measurements. Samples were stained for cell surface markers with antibodies conjugated with either FITC, PE or APC. For positive and negative controls, unstained and isotype control-treated cells were used, respectively. Further analysis was performed using FlowJo software (Tree Star, Ashland, OR).

### Antigen uptake, processing and phagocytosis assays

M1 and M2 macrophages (300,000/well, 96-well plate) were transfected on Day 7 with MALAT1 or control GapmeRs (Qiagen). MALAT1- or control GapmeR-transfected M1 and M2 Mφ were incubated with Texas Red-conjugated Ova and DQ-conjugated Ova (both 1 mg/ml, Molecular Probes, Grand Island, NY) in complete media to assess antigen uptake and antigen processing, respectively [29]. A phagocytosis assay was performed with pHrodo Red/Green-conjugated *E. coli* (Invitrogen, Carlsbad, CA) at 24 h post transfection, according to the manufacturer’s protocol [27].

### Macrophage bactericidal activity assay

To measure the bactericidal activity of macrophages, differentiated macrophages were incubated with *E. coli*, and the number of surviving bacteria was quantitated by MTS (3-(4,5-dimethylthiazol-2-yl)-5-(3-carboxymethoxyphenyl)-2-(4-sulfophenyl)-2H-tetrazolium)-based colorimetric assay [30–32]. CD14+ monocytes (50,000) were seeded per well of a 96-well plate and differentiated into M1 or M2 macrophages. LncRNA MALAT1 was silenced using ASO-MALAT1, and cells were incubated with E. coli (DH5α) particles. E. coli particles (from log phase cultures in LB media) were quantitated by measuring the O.D. at 600 nm (OD600 = 8.0X10^8^ cells/ml). Bacteria were washed and resuspended in DMEM, and 5X10^5^ particles were incubated per well of macrophages (MOI 10). Culture plates were centrifuged at 150×g for 5 min to bring the bacteria into contact with the macrophages. Plates were incubated at 37 °C for 8 h. After incubation, macrophages were lysed by Milli-Q water (hypotonic lysis) for 30 minutes. E. coli viability was determined using the CellTiter 96 AQueous Cell Proliferation Assay Kit (Promega, Madison, WI) according to the manufacturer’s protocol. At the same time, a bacterial titration standard containing a 2-fold dilution of bacteria in DMEM was prepared, and an MTS assay was performed. Absorbance from the bacteria titration plate was fitted to the standard curve from which viable bacteria numbers were calculated in ASO-MALAT1- or ASO-Control-transfected macrophages.

### Multiplex cytokine array

Cell culture supernatants (spent media) were collected from M1 and M2 macrophages challenged with *E. coli* LPS for 4 h and 24 h. The levels of 8 different cytokines (IL-1α, IL-1β, IL-6, IL-8, CXCL10, TNF-α, IL-1Rα and IL-10) were analyzed by multiplex assays using custom multiplex cytokine plates (EMD Millipore, Billerica, MA, USA). Data were collected on a MAGPIX instrument (Luminex, Austin, TX).

### LncRNA MALAT1 cloning

Bioinformatics prediction of miR-30b-5p and miR-26a binding sites in lncRNA MALAT1 was performed by using an online database, RNA hybrid (https://bibiserv.cebitec.uni-bielefeld.de/rnahybrid/). To perform a luciferase assay to test the predicted binding sites of miR-30b-5p and miR-26a within lncRNA MALAT1, a 1574 bp fragment of lncRNA MALAT1 (8-1582 bp) was PCR amplified (forward primer – TGCAGCCCGAGACTTCTGTAAAGG; reverse primer – GCTTCATCTCAACCTCCGTC) using genomic DNA from monocytes and ligated in dual luciferase reporter vector psiCHECK-2 downstream of Renilla luciferase (Promega Corporation) vector in Xho1 and Not1 sites in MCS. For overexpression of MALAT1 in completion luciferase assays with psiCHECK2-TLR8, the MALAT1 sequence was subcloned from psiCHECK-2 in pCDNA3.1 (pcDNA-MALAT1) within EcoRI and NotI sites.

### Dual Luciferase Assays

Dual luciferase reporter assays were performed using Promega dual luciferase kit, as previously described [27]. Briefly, HEK293 cells were seeded in 96-well plates, at a density of 3X10^4^ cells/well in DMEM supplemented with 10% fetal bovine serum. All transfections were performed using 0.5 μL of Lipofectamine 2000 (Invitrogen), 120 ng of dual luciferase reporter plasmid psiCHECK2 containing a lncRNA MALAT1 fragment having hsa-miR-30b binding site, and co-transfected with a final concentration of 10 nM, 25 nM and 50 nM of synthetic hsa-miR-30b or control mimics (50 nM) (Qiagen). At 36 h post-transfection, cells were lysed in passive lysis buffer (Promega Corporation), and dual luciferase assays were performed using the multilabel reader VictorTM x5 (Perkin Elmer 2030-Perkin Elmer Health Sciences Inc., Shelton, CT, USA).

### Transcriptome profiling

For transcriptome wide expression profiling microarray analysis was used. Dendritic cells were transfected with miR-30b-5p or control mimic and harvested after 36 h, and total RNA was isolated using the miRNeasy kit (Qiagen). We used Affymetrix HTA-2_0 arrays (Santa Clara, CA, USA) for gene expression profiling, and the analysis was performed as described previously [33]. Array data were in compliance with Minimum Information About a Microarray Experiment (MIAME) guidelines and deposited in the Gene Expression Omnibus public database under Accession Number GSE151262.

### Study population and sample collection

This study was conducted in accordance with the Declaration of Helsinki and approved by the Ethics Committee at the University Autónoma de Nuevo León Facultad de Odontología, Monterrey, Mexico, and Institutional Review Board at the University of Illinois at Chicago, College of Dentistry, Chicago, IL, USA (IRB # 2015-1093). Subjects presenting to the Postgraduate Periodontics Clinic at the Dental School of the Universidad Autonóma de Nuevo León were recruited for this study. Subjects (N=10) with chronic periodontal disease displayed probing depth ≥ 6mm with bleeding on probing and radiographic evidence of bone loss. Health periodontal patients displayed probing depths ≤ 3mm, with no bleeding on probing and no radiographic evidence of bone loss. Inclusion criteria included male and female patients ages 18 to 65 years and in good systemic health. Exclusion criteria included chronic disease (diabetes, hepatitis, renal failure, clotting disorders, HIV, etc), antibiotic therapy for any medical or dental condition within a month before screening, and subjects taking medications known to affect periodontal status (e.g. phenytoin, calcium channel blockers, cyclosporine). For periodontally healthy subjects (N=10), a single gingival biopsy sample (including gingival epithelium, col area, and underlying connective tissue) was collected at the time of crown-lengthening procedures. The biopsy sample was harvested using intrasulcular and inverse bevel incisions approximately 2 mm from the free gingival margin at the crest of the interproximal papillae extending horizontally capturing the interproximal col area and immediately placed in RNAlater (Qiagen, Gaithersburg, MD, USA) and stored at -80°C until further use.

### Murine model of ligature-induced periodontitis

Periodontitis was induced using a 6-0 silk ligature placed bilaterally between the maxillary first and second molar and local injection of *P. gingivalis* (2µl of 1X10^9^ pfu) around buccal and palatal gingiva) in 8-12-week-old female mice (n=3) under anesthesia using intraperitoneal injection of ketamine and xylazine. Animals were euthanized at 8 day post-ligature (DPL) placement. The mice with displaced ligature during experimental period were excluded from the evaluation. Gingival tissue were harvested from around the maxillary first and second molars and homogenized. Total RNA was extracted from the excised gingival tissue using the miRNeasy kit (Qiagen). RT-qPCR was performed as described above and the gene expression levels were normalized to those of the reference endogenous controls RNU6A and β-actin for miRNA and mRNA, respectively.

### Statistical Analysis

All of the data were analyzed and plotted using GraphPad Prism (La Jolla, USA). The results are represented as SD or SEM from three independent replicates. P values were calculated using Student’s t-test, and P < 0.05 was considered significant. *P < 0.05, **P < 0.01, ***P < 0.001.

## Results

### MALAT1 expression is responsive to macrophage differentiation and TLR4 stimulation

Our previous lncRNA PCR array profiling showed higher expression of MALAT1 in M2Mφ compared to monocytes, suggesting its role in macrophage differentiation [26]. Therefore, in this study, we first examined the expression dynamics of MALAT1 in monocytes during M1 and M2 Mφ differentiation at the following time points: 18 h, Day 3, Day 5, and Day 7. Our results showed a time-dependent upregulation of MALAT1 during both M1Mφ and M2Mφ differentiation (**Fig. 1A**). The expression of MALAT1 increased significantly at Day 3 and peaked at Day 5. Comparing the peaks of its expression at Day 5 with monocytes, we observed 7-fold and 3.7-fold increases in MALAT1 expression in M1 and M2 Mφ, respectively. These observations suggest that MALAT1 levels are induced during macrophage differentiation, however, its expression is particularly high in M1 subset.

**Figure 1.**
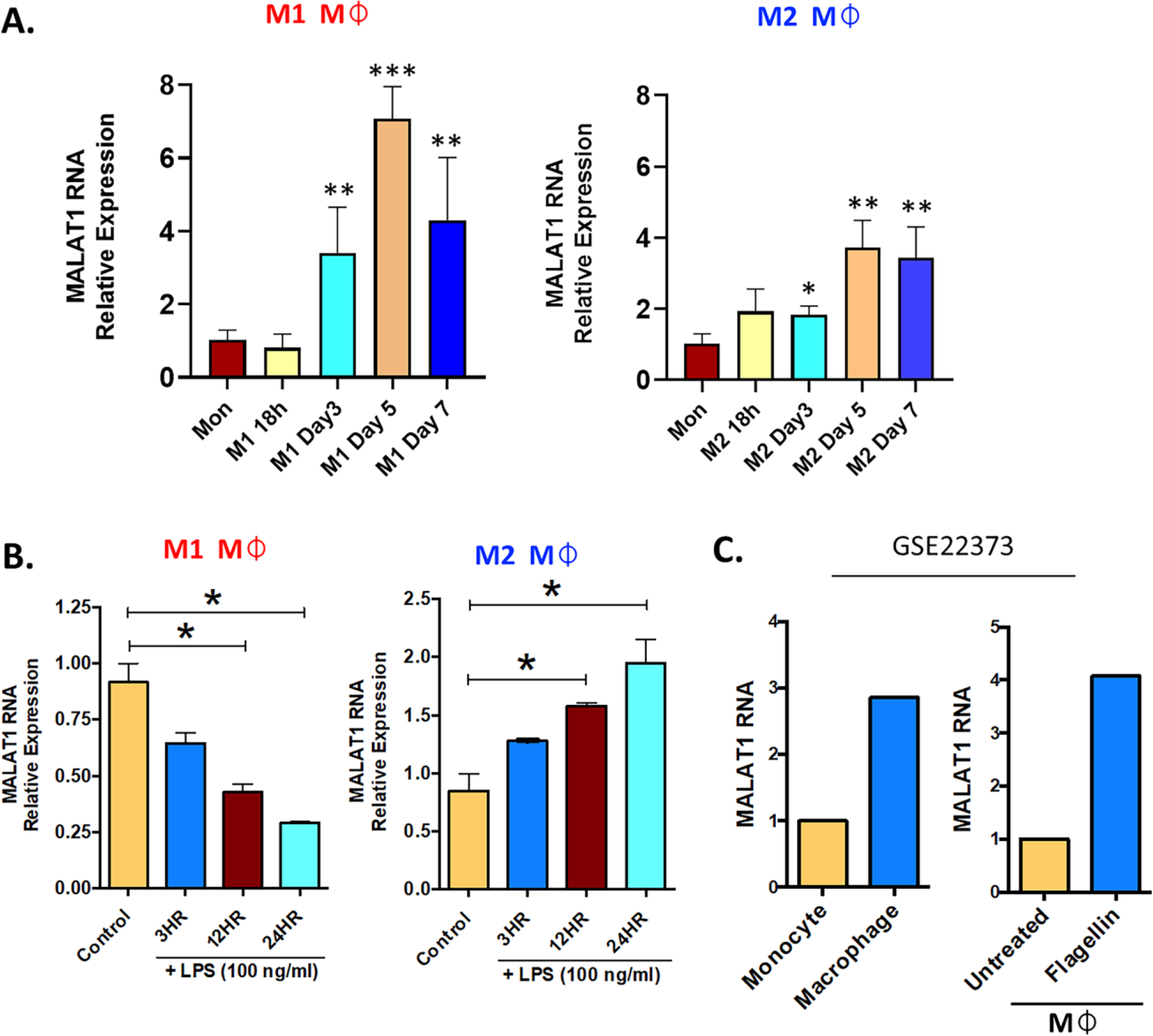
Long noncoding RNA MALAT1 is responsive to macrophage differentiation and activation *in vitro* and *in vivo*. (**A**) Expression of MALAT1 during the differentiation of M1 and M2 macrophages compared with monocytes *in vitro*. Primary CD14+ monocytes were isolated and differentiated into M1 Mφ or M2 Mφ in the presence of recombinant human (rh) GM-CSF (1000 U/ml) or M-CSF (50 ng/ml). Total RNA was isolated at various time points, and MALAT1 expression was quantified by RT–qPCR. (**B**) Time kinetics of MALAT1 expression in response to LPS stimulation (100 ng/ml) in M1 and M2 macrophages. Data (n=3) are representative of three independent experiments and are presented as the mean ± SEM. Significance was determined by Student’s two-tailed t-test. *P<0.05, **P<0.01, ***P<0.001. (**C**) *In vivo* macrophages express higher MALAT1 than monocytes and after TLR activation. MALAT1 RNA expression in a public dataset (GSE22373) in blood monocytes and intestinal macrophages or in macrophages treated with *S. typhimurium* flagellin (100 ng/ml) for 24 h.

Monocytes are also precursors to myeloid dendritic cells (mDCs). We then asked whether the induction of MALAT1 occurs during monocyte differentiation to DCs. MALAT1 expression was examined during DC differentiation and we observed that its levels significantly increase as early as 18 h (5-fold) and peaked at Day 3 (15-fold) of mDC differentiation (**Supplemental** **Fig. 1A**). Together, these observations clearly suggest that MALAT1 expression is induced during monocyte-to-Mφ and DC differentiation.

Macrophages are long-lived, TLR4+ myeloid cells, and dynamic transcriptional changes are integral to elicit adept immune responses. Therefore, we next evaluated whether MALAT1 is responsive to TLR4 ligation, viz. *E. coli* LPS challenge. M1 and M2 Mφ were treated with *E. coli* LPS (100 ng/ml) for 3, 12, and 24 h, and MALAT1 expression was quantified by RT–qPCR. In M2Mφ, MALAT1 was upregulated by LPS treatment as early as 3 h and was sustained until 24 h (2-fold increase) (**Fig. 1B**). In contrast, MALAT1 levels in M1 Mφ showed a progressive decrease in levels, suggesting that MALAT1 is differentially expressed in M1 and M2 Mφ in response to TLR4 activation and may contribute to polarization pathways (**Fig. 1B**).

Next, we asked whether MALAT1 expression is differentially regulated *in vivo*. For this, we examined MALAT1 expression in a published microarray dataset (GSE22373) of unstimulated monocytes and macrophages and flagellin-stimulated monocytes and macrophages. We noted higher expression of MALAT1 in macrophages than in monocytes and in response to flagellin treatment (TLR5 stimulation) (**Fig. 1C**; [34]).

Overall, these results show that MALAT1 expression is responsive to Mφ differentiation and activation and suggest that MALAT1 may regulate both aspects of Mφ biology.

### Knockdown of MALAT1 favors M2 macrophage phenotype

To elucidate the role of MALAT1 in Mφ polarization, we performed MALAT1 knockdown using antisense oligos (ASO-MALAT1). CD14+ monocytes were differentiated into M1 and M2 Mφ and then transfected with ASO-MALAT1 or ASO-Control at day 3. We observed significant MALAT1 silencing in ASO-MALAT1 transfected Mφ (both subtypes) compared to ASO-Control Mφ (**Supplemental** **Fig. 1B**). We did not observe any adverse effect of MALAT1 knockdown on the viability of M1 or M2 Mφ (**Supplemental** **Fig. 1C**) in the MTT assay.

Next, we examined the impact of MALAT1 knockdown on Mφ polarization by quantifying the expression of various M1 and M2 Mφ phenotype markers by RT–qPCR. Compared to the control, knockdown of MALAT1 resulted in increased mRNA expression of M2 markers ARG1 (4.8±0.87-fold), STAT3 (1.63±0.24-fold), CCL2 (4.69±1.19-fold), and IL-10 (1.45±0.1-fold) (**Fig. 2A; Upper panel**). In contrast, the expression of M1Mφ markers iNOS (1.68±0.87-fold), STAT1 (0.87±0.52-fold), TNF-α (1.34±0.49-fold), and ARG2 (1.81±0.73-fold) showed a trend of upregulation in MALAT1, albeit the changes were not significant (**Fig. 2A; lower panel**).

**Figure 2.**
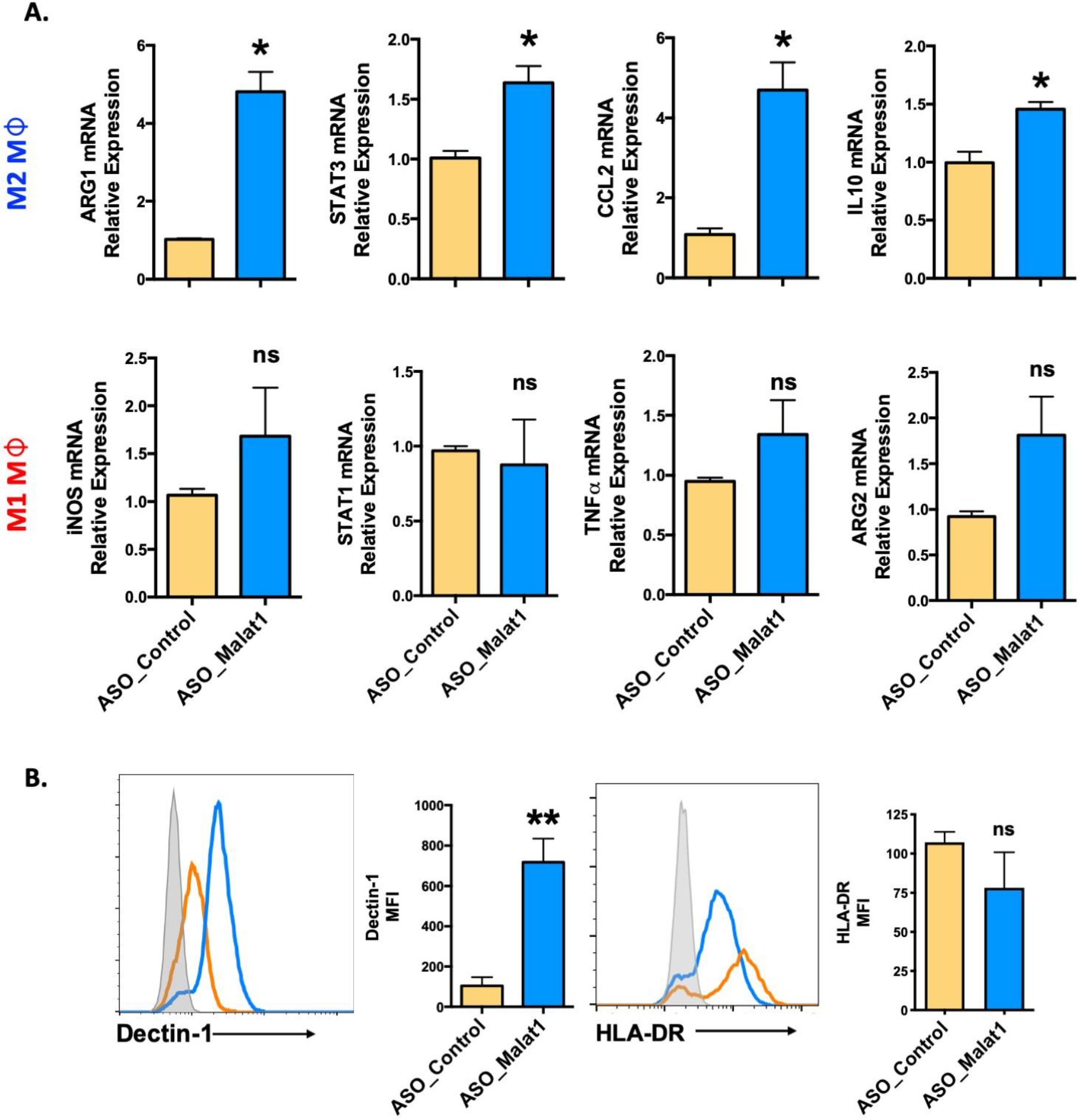
Knockdown of MALAT1 attenuates M1 and M2 macrophage differentiation from monocytes. (**A**) mRNA expression of macrophage polarization-specific genes in MALAT1 knockdown cultures. M1Mφ and M2Mφ were transfected with MALAT1 (ASO-MALAT1) or control (Control_ASO) GapmeR at a final concentration of 50 nM. After 48 h, total RNA was isolated to quantify M2 (ARG1, CCL2, STAT3, IL-10) and M1 (iNOS, STAT1, TNF-α, ARG2) markers by RT–qPCR using specific gene primers. GADPH was used as an internal control. The 2^-ΔΔCt^ method was used to calculate the expression compared to control GapmeR-transfected cells. (**B**) Histograms showing cell surface expression of M2 (Dectin-1) and M1 (HLA-DR) markers in MALAT1 knockdown macrophages. The gray shaded area represents isotype control, while orange and blue lines represent ASO-MALAT1 and ASO-Control, respectively. Percent geometric MFI values for Dectin-1 and HLA-DR in MALAT1- or control GapmeR-transfected cells. Data (n=4) are representative of three independent experiments and are presented as the mean ± SEM. Significance was determined by Student’s two-tailed t-test. *P < 0.05, **P < 0.01, ***P<0.001 (unpaired, two-tailed Student’s t-test), ns: non-significant.

To confirm the RT–qPCR results, cell surface markers for Mφ polarization, Dectin-1 (for M2Mφ), and HLA-DR (for M1Mφ), were evaluated by flow cytometry. **Supplemental** **Fig. 1D** shows the gating strategy for flow analysis. MALAT1 knockdown resulted in significantly higher (89.7±3.1%) Dectin-1+ population than the control (49.8±14.31%) (**Fig. 2B**). The mean fluorescence intensity (MFI) of Dectin-1 in M2Mφ was seven-fold higher (718%) in MALAT1 knockdown cells than in control transfected cells. Although HLA-DR+ cells slightly decreased in MALAT1 knockdown (73.7±20%) compared with ASO-control (100.33±11.96), this change was not statistically significant (**Fig. 2B**). Together, these experiments clearly suggest that MALAT1 downregulation leads to increased expression of M2 gene markers thereby favoring the M2 phenotype.

### MALAT1 regulates antigen uptake, processing, and phagocytosis in macrophages

Macrophages are classical antigen-presenting cells (APCs) that recognize diverse antigens, internalize them by endocytosis or phagocytosis, process them, and finally present on their cell surface for T cell activation [35]. To study the effect of MALAT1 knockdown on the function of macrophages, we evaluated antigen uptake, processing and bacterial phagocytosis in MALAT1 knockdown M2Mφ. To examine the effect of MALAT1 knockdown on antigen uptake and processing, Mφ were treated with Texas red-labeled ovalbumin and DQ-ovalbumin, respectively. DQ-ovalbumin is labeled with BODIPY dye with a fluorogenic quencher, which is released upon hydrolysis by cellular proteases. Extracellular DQ-ovalbumin is non-fluorogenic and becomes fluorogenic upon uptake and processing by macrophages.

Knockdown of MALAT1 in M2Mφ resulted in decreased antigen uptake (**Fig. 3A**). The percentage of Texas red-positive cells significantly decreased from 100.4±7.8% in ASO-Control cells to 67.7±5.4% in MALAT1 knockdown cells (**Fig. 3B; left panel**). Geometric MFI values showed ∼30% reduction in florescence intensity further showing attenuation of antigen uptake (**Fig. 3B; right panel**). The processing of internalized ovalbumin in MALAT1 knockdown Mφ was also decreased, as observed by imaging (**Fig. 3C**). Similar results were observed in flow cytometry analysis (**Fig. 3D**). The percentage of BODIPY-positive cells and the MFI were significantly decreased (∼40%) in MALAT1 knockdown compared to control cells.

**Figure 3.**
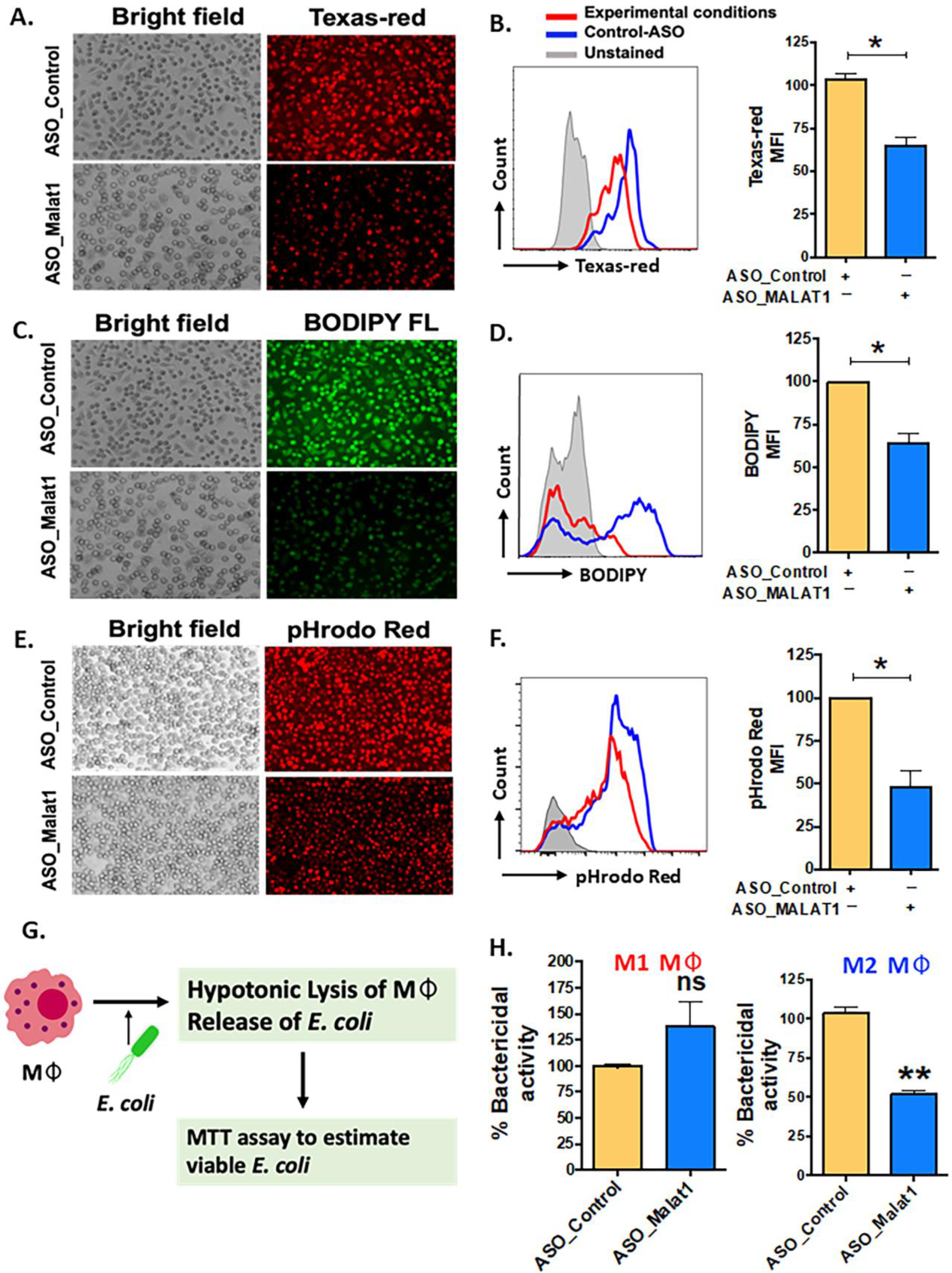
MALAT1 knockdown regulates antigen uptake and processing by macrophages. Cells were transfected with ASO-MALAT1 or ASO_Control oligos and after 48 h incubated with Texas Red-conjugated Ova (for antigen uptake), BODIPY-Ova (for antigen processing) or pHrodo™ Red *E. coli* BioParticles™ (Invitrogen) for 2-4 hours. Representative florescent microscopy images showing the inhibitory impact of MALAT1 knockdown on (**A**) antigen uptake, (**C**) antigen processing or (**E**) phagocytosis. In addition, Mφ were harvested after incubation with different fluorescent probes and analyzed by flow cytometry. Overlay of representative histograms of ASO_MALAT1- or ASO_Control-transfected cells showing differences in (**B**) Texas red, (**D**) BODIPY and (**F**) rhodamine florescent signals for each of the biological assays examined. Bar graphs showing corresponding percent geo. MFI values next to each histogram. The data are presented as the ±SEM of four independent experiments in each cell type. Student’s t-test was conducted to calculate p values. **P < 0.01, ***P < 0.001. (**G**) Schematic showing the workflow of the macrophage bactericidal activity assay. (**H**) Bactericidal activity assay in MALAT1 knockdown M1 and M2 macrophages. In, B, D, H, F: Each bar shows the average±SD of three independent experiments (n=3). \**P*<0.05, \*\**P*<0.01, and \*\*\**P*<0.001 (unpaired, two-tailed Student’s t-test).

To assess the effect of MALAT1 knockdown on phagocytosis, cells were treated with pHrodo-Red-labeled *E. coli,* and bacterial uptake was quantified by imaging and flow cytometry analysis. Knockdown of MALAT1 remarkably attenuated bacterial phagocytosis by M2Mφ, as shown in **Fig. 3E**. Microscopy results were independently verified by flow cytometry (**Fig. 3F**), as observed by a significantly lower percentage of pHrodo red-positive cells. Quantitation of geometric MFI also showed reduced uptake in MALAT1 knockdown (48.13%) cells compared to control (set at 100).

Bactericidal activity is a characteristic function of M1Mφ. We next asked whether MALAT1-mediated skewing of the Mφ phenotype toward M2 could attenuate bacterial killing potential. Cells were transfected with ASO-MALAT1 or ASO-Control and challenged with live *E. coli* at different concentrations. **Fig. 3G** shows the workflow of the macrophage bactericidal activity assay. To quantitate the bacterial counts, we standardized the absorbance of *E. coli* culture in the MTS assay to plot a standard curve for calculating the unknown bacterial numbers and used this curve to assess bactericidal activity (**Supplemental** **Fig. 2A**). MALAT1 knockdown in M2Mφ showed significantly less bactericidal activity (51.76%) than that in ASO-Control transfected M2Mφ (set at 100%) (**Fig. 3H**). Higher numbers of viable *E. coli* (4.4X10^6^) were observed in MALAT1 knockdown M2Mφ compared to the control (2.31X10^6^), suggesting that lower MALAT1 expression impairs bactericidal activity (**Supplemental** **Fig. 2B**). In contrast, MALAT1 knockdown in M1Mφ showed no statistical differences in bactericidal activity, as observed by similar levels of viable *E. coli* counts (2.14X10^6^) compared to the control (3.09X10^6^) (**Fig. 3H**; **Supplemental** **Fig. 2B**). These results suggest that the knockdown of MALAT1 attenuates phagocytosis, antigen processing and differentially impairs bactericidal activity of M2Mφ.

### MALAT1 knockdown leads to decreased inflammatory cytokine secretion

Cytokine secretion is tightly coupled with endocytosis or phagocytosis [36, 37]. Having established that MALAT1 knockdown negatively regulates phagocytosis, we next asked whether it also affects the innate immune response by modulating the cytokine secretion. M1 or M2Mφ were transfected with ASO-MALAT1 or ASO-Control for 48 h and then challenged with *E. coli* LPS (100 ng/ml). Supernatants collected at 4 and 24 h were examined for secreted levels of eight different cytokines/chemokines, including proinflammatory cytokines (IL-1α, IL-1β, IL-6, IL-8, CXCL10, and TNF-α) and anti-inflammatory cytokines (IL-1Rα and IL-10). Most of the cytokines examined (except IL-8) showed significant changes in expression either at an early or late time point in MALAT1 knockdown cells compared to control cells, suggesting that MALAT1 regulates innate immune responses in macrophages.

Compared to ASO-Control macrophages, MALAT1 knockdown M1 and M2 macrophages showed significant downregulation in the levels of IL-6, TNF-α, IL-1Rα and IL-10, while IL-1α, IL-1β, and CXCL10 exhibited antagonistic expression in M1 and M2 macrophages (**Fig. 4A**). Downregulation of the proinflammatory cytokines IL-6 and TNF-α was observed at both 4 and 24 h in M2 macrophages, but a significant reduction was observed at 4 h in M1 macrophages. The anti-inflammatory cytokines IL-10 and IL-1Rα displayed strikingly similar downregulation in both M1 and M2 macrophages at both the time points examined. These results indicate that MALAT1 knockdown has a similar impact on anti-inflammatory cytokines in both M1 and M2 macrophages but causes differential changes in proinflammatory cytokine levels. No significant changes were noticed for IL-8 expression in MALAT1 knockdown cells. **Fig. 4B** **& C** summarizes the modulation of various cytokines in MALAT1 knockdown cells.

**Figure 4.**
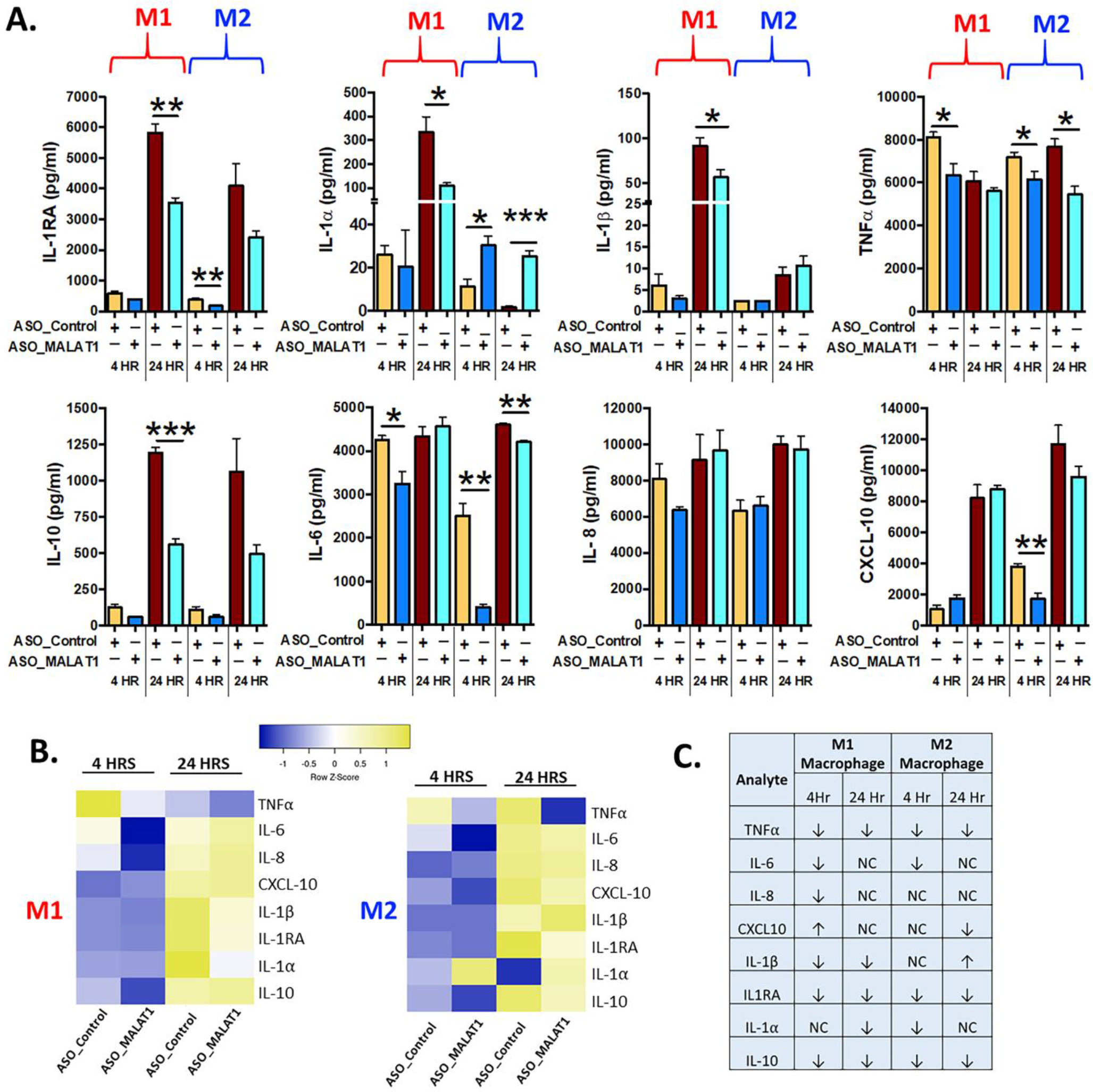
MALAT1 knockdown regulates inflammatory cytokine secretion by macrophages. Inflammatory cytokines were measured in the culture media of MALAT1-depleted M1 and M2 macrophages stimulated with *E. coli* LPS for 4 and 24 hours. **(A)** Bar graph showing proinflammatory (IL-1Rα, IL-1β, IL-1β, TNF-α) and anti-inflammatory cytokine (IL-10, IL-6, IL-8, CXCL10) levels in M1 and M2 macrophages (n=4). \**P* < 0.05, \*\**P* < 0.01, and \*\*\**P* < 0.001 (unpaired, two-tailed Student’s t-test). **(B)** Heatmap of pro- and anti-inflammatory cytokines in M1 and M2 macrophages. **(C)** Table summarizing the expression of pro- and anti-inflammatory cytokines in M1 and M2 macrophages.

Interestingly, MALAT1 knockdown reduced the levels of IL-1α and IL-1β at the 24 h time point in M1 Mφ and caused upregulation of IL-1α (both at 4 and 24 h) in M2 Mφ, but no significant changes were observed for IL-1β in M2 macrophages (**Fig. 4A**). Similarly, the proinflammatory chemokine CXCL10 was differentially impacted by MALAT1 knockdown in M1 and M2 Mφ. Reduced levels of CXCL10 were observed in M2Mφ at both time points; although it was significant at 4 h, no significant impact on CXCL10 levels was observed in M1Mφ (**Fig. 4A**). These findings validate our previous observations that MALAT1 displays distinct functionality in M1 and M2 macrophages. Overall, cytokine profiling in *E. coli* LPS-challenged M1 and M2 macrophages strongly supports that MALAT1 controls the innate immune response by supporting the proinflammatory immune response and that MALAT1 knockdown has a differential impact on M1 and M2 macrophages.

### MALAT1 acts as an endogenous sponge of miR-30b and promotes the expression of its targets

Recent studies have shown that lncRNAs can bind and sequester mature miRNAs [16,24,25]. By regulating the bioavailability of functional miRNAs, lncRNAs can modulate the expression of their target protein-coding genes. Our laboratory has previously reported a key association between the expression dynamics of miRNAs and M-CSF-mediated monocyte-to-Mφ differentiation [28]. miR-30 family members were among the significantly downregulated miRNAs, and we demonstrated their role in potentiating the differentiation and function of both macrophages and DCs [27, 28]. The expression of all five members of the miR-30 family (miR-30a-e) was significantly downregulated during M2 macrophage differentiation (**Supplemental** **Fig. 2C**) and exhibit an antagonistic relationship with MALAT1 levels (**Fig. 1A**). Interestingly, compared to M1Mφ, miR-30b is expressed at higher levels in M2Mφ suggesting a differential role of miR-30b in macrophage polarization (**Fig. 5A**). Therefore, we hypothesized that overexpression of miR-30 family members would antagonize MALAT1 function by promoting the M2 phenotype.

**Figure 5.**
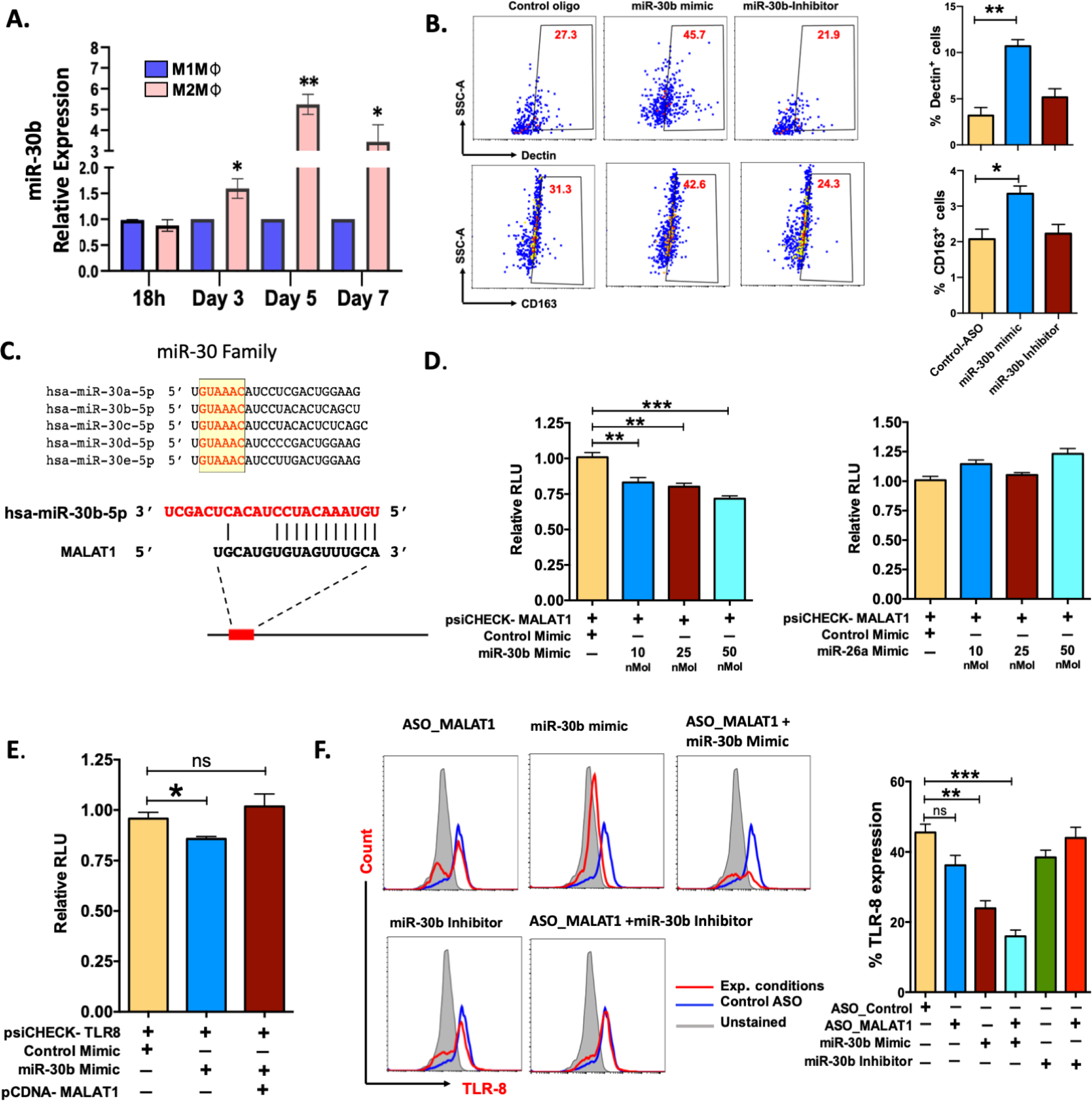
MALAT1 acts as an endogenous sponge of miR-30b and promotes the expression of its targets. **(A)** Comparative miR-30b expression levels in differentiating M1 and M2 macrophages. Total RNA isolated from monocyte-derived M1 and M2 Mφ were examined for miR-30b expression by RT-qPCR. Histograms showing M1 normalized miR-30b expression in M2 Mφ. RNU6B was used a endogenous control. **(B)** miR-30b overexpression promotes the M2 phenotype. Day 4 M2Mφ were transfected with miR-30b mimic, inhibitor, or control mimic (50 nM), and M2-specific markers, viz. Dectin-1 and CD163 expression, were assessed by flow cytometry. Representative dot plots show the percent cell population positive for the markers. Bar graphs (on the right side of dot plot) show overall Dectin-1+ and CD163+ cells in two independent experiments (n=3). (**C**) Sequence alignment of predicted complementary interactions of miR-30 family members and MALAT1. Conserved seed sequence is shown in yellow box. (**D**) Functional miR-30b interaction with MALAT1. The dual luciferase reporter plasmid psiCHECK2 (120 ng) cloned with the lncRNA MALAT1 fragment containing the predicted hsa-miR-30b binding site was cotransfected with various concentrations of hsa-miR-30b (10, 25 or 50 nM concentrations) or control mimic (50 nM) in HEK-293T cells. miR-26a, another miRNA with a binding site on MALAT1, was also screened by dual luciferase assay. At 36 h posttransfection, cells were lysed to quantify Renilla and firefly luciferase activity. Renilla activity was normalized to firefly activity, and the ratios subsequently normalized to vector transfected with control mimic set as 1. Data are expressed as the ±SEM of four independent transfections. (**E**) Competitive dual luciferase assay with psiCHECK2-TLR8 cotransfected with miR-30b mimic alone or in conjunction with pcDNA-MALAT1 with the miR-30b binding site. Bar graphs show reduced normalized Renilla activity in miR-30b cells, which was restored in pcDNA-MALAT1-transfected cells. Data are expressed as the ±SEM of four independent transfections. (**F**) MALAT1 functionally titrates miR-30b levels. Cells were transfected with ASO_Control, ASO_MALAT1, miR-30b mimic, miR-30b with ASO_MALAT1, and miR-30b inhibitor with ASO_MALAT1, and intracellular expression of TLR8 was examined by flow cytometry. Representative overlay histograms show TLR8 expression in different combinations, supporting that MALAT1 acts as an endogenous competitor of miR-30b. Bar graphs with the mean percent TLR8+ population are plotted from the same experiments. Each bar shows ±SD of 3 independent experiments (n=3). *P<0.05, **P<0.01, and ***P<0.001 (unpaired, two-tailed Student’s t-test).

To test this hypothesis, Day 4 differentiated M2Mφ were transfected with miR-30b mimic, inhibitor, or control mimic, and M2 markers (Dectin-1 and CD163) were quantified on Day 7. Flow cytometric analysis showed that M2Mφ transfected with miR-30b mimic upregulated both Dectin-1 and CD163. We observed a significantly higher percentage of Dectin-1+ cells in miR-30b overexpressing M2Mφ (44.9±4.75%; *P<0.005*) than in the control (23.7±3.55%) (**Fig. 5B**). Conversely, suppression of miR-30b by an inhibitor reduced Dectin-1 expression (21.1±1.5%) compared to miR-30b mimic (**Fig. 5B**). Similarly, the expression of CD163 was upregulated in miR-30b mimic-transfected cells (48.3 ±6.64%; *P<0.022*) compared to control (32.0±3.21%) or inhibitor-transfected cells (24.5±4.55%). These data suggest that miR-30b expression in monocytes favors the M2 phenotype rather than the M1 phenotype and functionally antagonizes the role of MALAT1 in shaping macrophage polarization (**Fig. 5B**).

In an independent transcriptome analysis of monocyte-derived dendritic cells overexpressing the miR-30b mimic, we identified various genes regulated by miR-30b involved in antigen processing and processing, phagosome, and endosomal pathways (most enriched GO terms) (**Supplemental** **Fig. 3A-D**). We bioinformatically predicted the upstream regulators viz. transcription factors that regulate differentially expressed genes in miR-30b overexpressing dendritic cells. Network analysis based on the transcription factors revealed that monocyte differentiation, M1 polarization, and M2 polarization pathways (among other signaling pathways) were enriched in miR-30b transfected dendritic cells suggesting that miR-30b exhibit a propensity to target gene networks involved in macrophage polarization **(Supplemental** **Fig. 3E, F)**. Although these analyses were performed in CD14+ monocyte-derived dendritic cells, they indicate a transcriptome-wide role of miR-30b in macrophage differentiation and polarization.

Antagonistic expression and function of miR-30b and MALAT1 prompted us to ask whether MALAT1 negatively regulates mature miR-30b levels. Detailed bioinformatics analysis was performed to scan potential miRNA binding sites within the MALAT1 sequence, and we identified a novel sequence complementary to the miR-30 family seed region (**Fig. 5C; Supplemental** **Fig. 2D**). To substantiate our *in silico* analysis, we cloned a partial MALAT1 sequence harboring the hsa-miR-30b recognition site into the psiCHECK2 vector for a dual luciferase assay. HEK293T cells were transfected with psiCHECK2-MALAT1 and miRNA or control mimic (at final concentrations of 10, 25, and 50 nM), and dual luciferase assays were performed after 36 h. Compared to the control mimic, we observed a dose-dependent reduction in Renilla luciferase activity in MALAT1 and miR-30b mimic cotransfected cells, confirming the presence of a functional miR-30b binding site on MALAT1 (**Fig. 5D; left panel**). Luciferase assays were also performed with psiCHECK2-MALAT1 and miR-26a, another miRNA similar to miR-30b that is downregulated during M2Mφ differentiation (data not shown) and predicted to bind MALAT1 in our bioinformatic screening (**Supplemental** **Fig. 2D**). However, we did not observe significant changes in luciferase activity compared to the control (**Fig. 5D; right panel**). These results confirm that MALAT1 physically interacts with miR-30b via a novel interacting site and may affect its biological functions.

### MALAT1 relieves miR-30b gene targets from posttranscriptional suppression

Next we asked whether MALAT1 promotes miR-30b target expression by titrating its levels. Studies from our laboratory have identified TLR8 and FcεRIγ as novel targets of miR-30b [38; *Naqvi et al., unpublished results*]. To test whether MALAT1 can functionally sequester miR-30b and relieve its cognate targets, we performed a luciferase assay with psiCHECK2-TLR8 cotransfected with miR-30b mimic and MALAT1 overexpressing plasmid pCDNA-MALAT1. Transfection of miR-30b with psiCHECK2-TLR8 reduced renilla luciferase activity (∼20%; *P*<0.01), while the presence of MALAT1 inhibited the miR-30b mimic-mediated reduction in luciferase activity by sequestering miR-30b, as evident from comparable renilla luciferase activity (no significant change) to psiCHECK2-TLR8 cotransfected with control mimic (**Fig. 5E**).

To confirm the luciferase assays, we examined the surface expression of TLR8 and FcεRIγ in MALAT1 knockdown, miR-30b overexpression, miR-30b inhibition, or a combination of these treatments by flow cytometric analysis. Transfection of the miR-30b mimic simulated the overexpression scenario, which could not be sequestered by the physiological levels of MALAT1; thus, we observed significantly low TLR8 expression (23.3±3.3%) compared with the control mimic (45.5±4.0%; *P<0.005*) (**Fig. 5F**). Conversely, we observed a decrease in TLR8 expression in MALAT1 knockdown, further supporting an inverse relationship between these noncoding RNAs. Flow cytometry data showed reduced TLR8 expression (36.2±4.9%; *P=0.079*) in MALAT1 knockdown cells compared to control cells (45.5±4.0%). This could be attributed to the reversal of MALAT1-mediated sequestering of miR-30b (**Fig. 5F**). To validate this further, we transfected cells with ASO-MALAT1 and miR-30b mimic and observed a highly significant reduction in TLR8 (*P<0.001*). The observed TLR8-expressing cells (15.9±3.11%) were significantly less abundant than miR-30b or MALAT1 alone (**Fig. 5F**). Transfection of miR-30b inhibitor with or without ASO-MALAT1 resulted in 38.4±3.5% and 43.9±5.2% TLR8+ populations, respectively, suggesting that inhibition of miR-30b restored TLR8 expression. However, miR-30b inhibition showed more potent restoration without MALAT1 knockdown, further substantiating the functional sequestration of miR-30b by MALAT1.

Furthermore, we evaluated the expression dynamics of FcεRIγ, a previously identified target of miR-30b from our laboratory [38]. FcεRIγ levels showed similar expression profiles as those observed for TLR8 under different transfection conditions. The percentages of FcεRIγ-expressing cells were 39.26 (±3.13), 19.3% (±0.98), 15.0% (±1.4), 11.4% (±2.26), 26.7% (±3.77), and 35.1% (±1.60) in cells transfected with ASO-Control, ASO-MALAT1, miR-30b mimic, ASO-MALAT1 + miR-30b mimic, miR-30b inhibitor and ASO-MALAT1+miR-30b inhibitor, respectively (**Supplemental** **Fig. 4A,B**).

Using two validated miR-30b targets, we show that MALAT1 acts as an endogenous sponge, and thus, its expression levels may play a critical role in M2Mφ phenotype by regulating miR-30 levels.

### MALAT1/miR-30b axis regulates phagocytosis and antigen processing in macrophages

After establishing that miR-30b sequestration by MALAT1 relieves miRNA-mediated repression of its target messenger RNAs (mRNAs), we next asked whether perturbing the MALAT1/miR-30b axis can impair Mφ innate functions viz. phagocytosis and antigen processing, which eventually bridge the innate and adaptive arms of immunity [39]. To test this hypothesis, M2Mφ were transfected with ASO-Control, ASO-MALAT1, miR-30b mimic, ASO-MALAT1 + miR-30b mimic, miR-30b inhibitor, and ASO-MALAT1 + miR-30b inhibitor, and the phagocytosis of rhodamine-labeled *E. coli* and processing of BODIPY-conjugated soluble antigen ovalbumin (Ova) were assayed. Compared to the control, a significant decrease in bacterial phagocytosis was noted in cells treated with the miR-30b mimic (**Fig. 6A**). We observed a significant attenuation of bacterial uptake in M2Mφ treated with ASO-MALAT1, as shown before (**Fig. 6A**; **Fig. 3**). Flow cytometry analysis showed a significant decrease in bacterial phagocytosis in MALAT1 knockdown (37.0±4.3%) and miR-30b overexpression (21.0 ±4.0%) compared to the control (74.0±4.35%) (**Fig. 6B**). These results clearly demonstrate that lncRNA MALAT1 and miR-30b regulate phagocytosis in M2 Mφ in an antagonistic manner. Both fluorescence microscopy and flow cytometry data showed significantly attenuated bacterial uptake in MALAT1 knockdown and miR-30b-overexpressing cells. Our flow cytometry data showed further decrease (15.3±4.50%) in pHrodo positive cells in ASO-MALAT1 and miR-30b mimic cotransfection, a significant reduction compared to either MALAT1_ASO (37.0±4.3%) or miR-30b overexpression (21.0 ±4.0%) alone. To further validate that the above effect on phagocytosis is driven by the MALAT1 and miR-30b interaction, we transfected M2Mφ with miR-30b inhibitor alone or with ASO-MALAT1 and observed a significant reversal of the miR-30b effect. The observed values of bacterial phagocytosis were 61.3.0±5.1% and 71.3±8.62% for the miR-30b inhibitor and miR-30b inhibitor + ASO-MALAT1 groups, respectively (**Fig. 6B**). These results substantiate the deterministic role of MALAT1 and miR-30b in phagocytosis.

**Figure 6.**
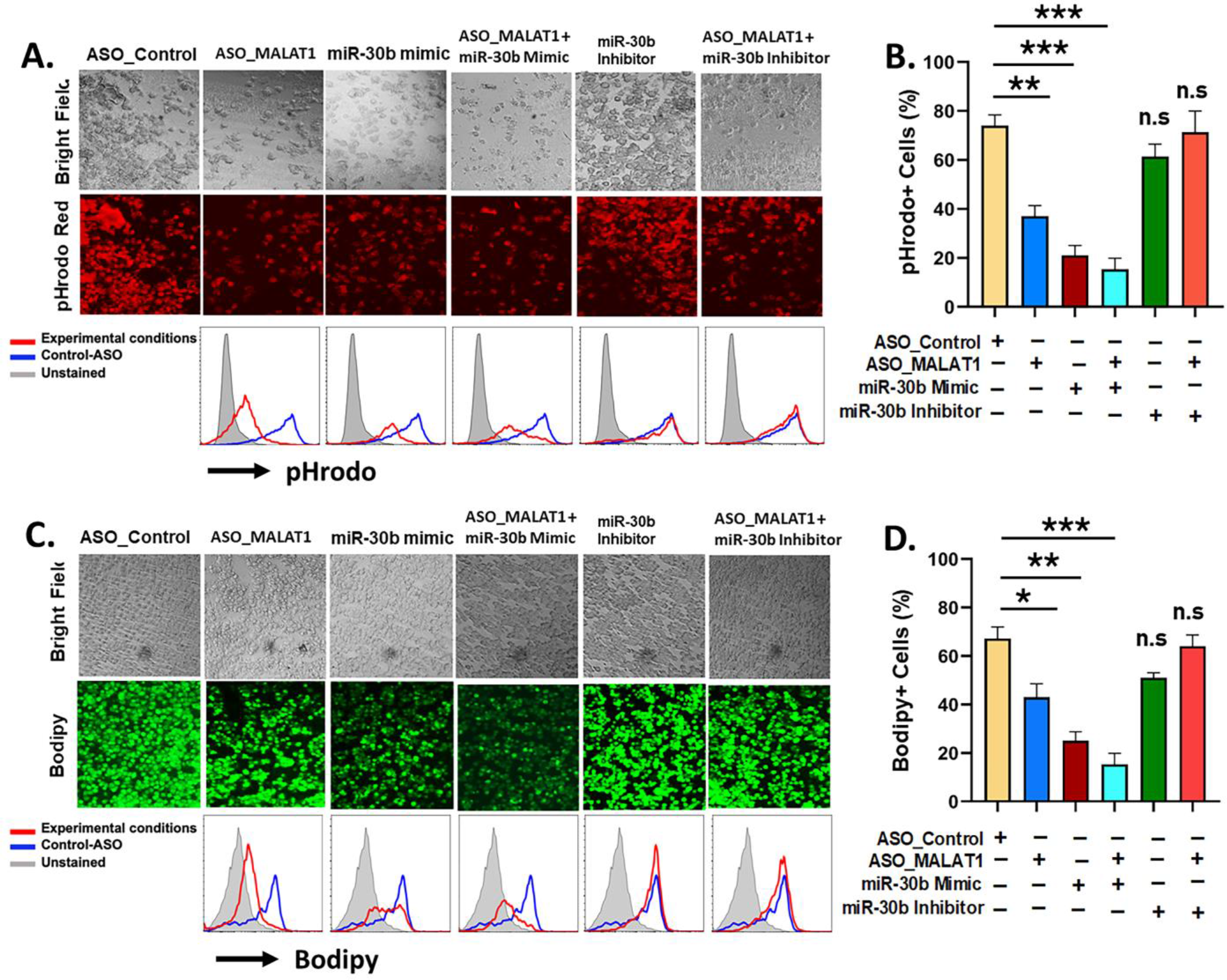
MALAT1 and miR-30b exhibit antagonistic regulation of phagocytosis and antigen processing in macrophages. (**A**) Phagocytosis assays were performed with labeled *E. coli* in Mφ transfected with ASO_Control, ASO_MALAT1, miR-30b mimic, miR-30b with ASO_MALAT1, and miR-30b inhibitor with ASO_MALAT1. Representative images were captured using a fluorescence microscope and quantified by flow cytometry, as shown by overlay histograms. (**B**) Bar graphs showing the rhodamine+ population analyzed by flow cytometry. (**C**) Same as (A) but cells were assessed for antigen processing after incubation with BODIPY-Ova. BODIPY release (green signal) indicating antigen processing was captured using a fluorescence microscope and quantified by flow cytometry, as shown by overlay histograms. (**D**) Bar graphs showing the BODIPY+ population analyzed by flow cytometry. The data presented are representative of three independent experiments (n=3). *P<0.05, **P<0.01, and ***P<0.001 (unpaired, two-tailed Student’s t-test), n.s: non-significant.

Since our above data strongly determined the effect of the MALAT1/miR-30b axis on phagocytosis, we next addressed whether the MALAT1 and miR-30b axes regulate antigen processing in M2 Mφ. To do so, the role of MALAT1 along with miR-30b in antigen processing of Ova was evaluated. M2Mφ transfected with ASO-MALAT1 showed a remarkable decrease in BODIPY (green signal) fluorescence under the microscope (**Fig. 6C**). These data were further validated by flow cytometry. Herein, we observed a significant (*P*<0.005) reduction in percent antigen processing in M2Mφ treated with ASO-MALAT1 (43.0.0±5.56%) compared to ASO-Control (67.2±4.7%). miR-30b mimic treatment exhibited more robust suppression of antigen processing, as observed by 25.3±3.6% BODIPY-positive cells (**Fig. 6D**). The reduction in antigen processing further in ASO-MALAT1- and miR-30 mimic-treated M2Mφ strongly suggests an antagonistic role in antigen processing. These data were further validated by reversal of antigen processing suppression upon treatment with a miR-30 inhibitor, either alone or in conjunction with ASO-MALAT1. The observed values of antigen processing in M2Mφ treated with miR-30b inhibitor alone or with MALAT1 were 51.2.3±2.05% and 64.0±4.68%, respectively, approaching the value of ASO-Control-treated cells (**Fig. 6D**). Together, these results clearly demonstrate that the MALAT1/miR30b axis in macrophages is functionally important in regulating innate immune functions such as antigen processing and phagocytosis.

### Higher MALAT1 expression correlates with reduced miR-30b levels and increased proinflammatory macrophage phenotype markers in inflamed gingival tissues

Our *in vitro* studies suggest that MALAT1 favors M1 phenotype by suppressing M2 promoting miR-30b. To confirm these findings, we examined MALAT1, and miR-30b expression in gingival biopsies collected from periodontally healthy and diseased human subjects as well as mice with ligature-induced periodontitis. Compared to healthy subjects, inflamed gingival biopsies showed markedly higher levels of MALAT1 (Fold change) and reduced expression of miR-30b (***Fig. 7A, B*)**. These results corroborate with the murine model of periodontitis, where we observed significantly higher MALAT1 and downregulation of miR-30b at 8DPL, which marks disease establishment and simulates human periodontitis subjects (***Fig. 7C, D***). These findings strongly suggest *in vivo* functional antagonism of MALAT1 and miR-30b.

**Figure 7.**
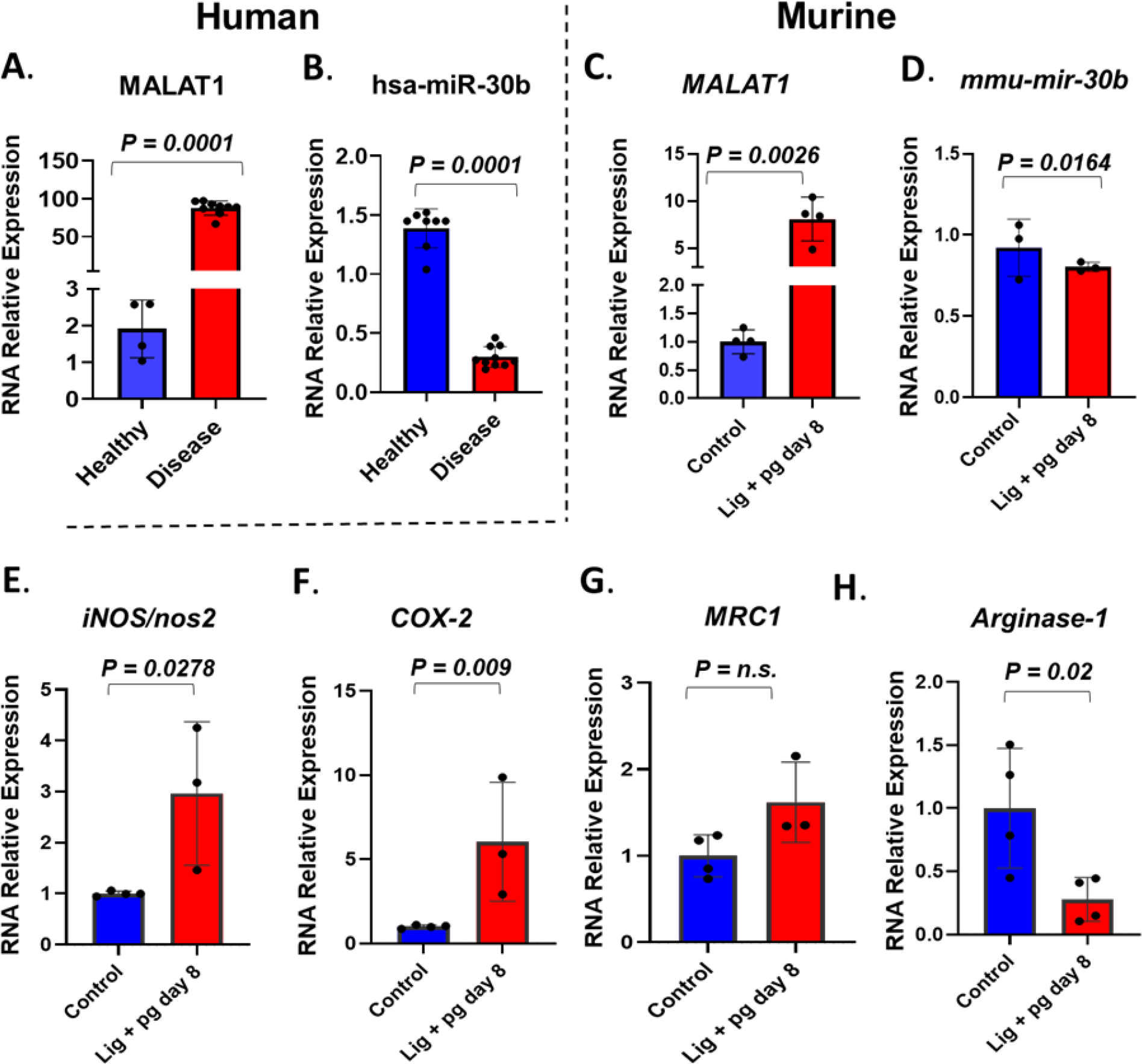
MALAT1/miR-30b exhibit antagonistic expression in inflamed human and murine gingival biopsies. Total RNA was isolated from healthy and inflamed gingival biopsies from human (n=10/group) and mice (n=3-4/group) subjected to ligature-induced periodontitis and the expression of MALAT1 and miR-30b was examined by RT-qPCR. Histograms showing relative fold change expression of (**A, C**) MALAT1 and (**B, D**) miR-30b-5p in periodontally healthy and inflamed human and murine gingiva. Transcript expression was normalized to β-actin (for MALAT1) and small nuclear RNA, RNU6B (for miR-30b). Expression of M1 macrophage phenotype markers is induced in inflamed murine gingiva. Histograms showing normalized expression of (**E**) *inos*, (**F**) *cox2* (both M1 markers), (**G**) *MRC1*, (**H**) *arginase1* (both M2 markers) in murine gingiva. β-actin was used as endogenous control. Data is presented as mean ± SEM. Student’s t-test was used to calculate P values.

We next asked whether higher MALAT1 and lower miR-30b affect macrophage polarization *in vivo*. Studies from other labs and ours (*Uttamani et al., unpublished results*) have demonstrated abundance of M1 phenotype macrophages in inflamed human gingiva [40]. We therefore examined M1 and M2 markers in healthy and inflamed murine gingival biopsies. Our results show significantly higher expression of *inos2* and *cox-2* compared to animals without ligature (***Fig. 7E, F***). Expression of *arginase 1*, an M2 marker, was significantly downregulated in inflamed gingiva, while *MRC1* levels did not show significant change (***Fig. 7 G,H***). Overall, these results strongly support that higher MALAT1 expression correlates with M1 markers and exhibit antagonism with miR-30b *in vivo* indicating a functional role of MALAT1 in shaping macrophage polarization.

## Discussion

Macrophage polarization is a highly coordinated biological process that involves the integration of numerous factors acting at the transcriptional, posttranscriptional, and posttranslational levels. Noncoding RNAs (ncRNAs) have been demonstrated to play deterministic roles in macrophage polarization. Despite a large number of known lncRNA transcripts, only a few are characterized for their functional role in macrophage polarization. This study evaluated lncRNA MALAT1 in macrophage polarization and studied its crosstalk with miR-30b, via a novel interaction site. We demonstrated the biological significance of this interaction in periodontally diseased human and murine gingival biopsies.

The relationship between MALAT1 expression and macrophage phenotype is poorly understood and needs further investigation. MALAT1 expression analysis in M1 and M2 Mφ exhibit differential response suggesting a unique functional requirement of this lncRNA by polarized Mφ. We therefore focused on delineating whether MALAT1 favors a specific macrophage phenotype. Knockdown of MALAT1 in M2Mφ significantly induced the expression of multiple M2 markers, including Arg1, STAT3, CCL2, IL-10, and Dectin-1, and concomitantly downregulated M1 markers (STAT1, CXCL10, HLA-DR), albeit not significantly. In line with our results, Cui et al. [41] reported that MALAT1 knockdown in IL-4-generated murine M2Mφ promoted the M2 phenotypic markers arginase 1 (Arg-1), YM-1, and mannose receptor C-type 1 (MRC1). Moreover, alveolar macrophages from MALAT1-deficient mice display the M2 phenotype, supporting a proinflammatory function of MALAT1. Consistent with our observations, knockdown of MALAT1 suppresses proinflammatory cytokine secretion, favoring the M1 phenotype [42], while delivery of MALAT1 through extracellular vesicles promotes the M1 phenotype via upregulation of HMGB1 [43]. In contrast, Huang et al. [44] demonstrated that exosomes from oxidized low-density lipoprotein (oxLDL)-treated endothelial cells deliver MALAT1 to monocytes and promote the M2 phenotype [44]. MALAT1 knockdown in hepatocellular carcinoma cells inhibits VEGF-A levels and skews macrophages toward the M1 subtype [45]. Thus, Mφ polarization is differentially responsive to endogenous and exogenous factors and MALAT1 is one of the critical molecular determinants in the process. Overall, these observations support an unequivocal role of MALAT1 in macrophage polarization,

The functional effect of MALAT1 on macrophage innate immune functions, viz. phagocytosis and antigen presentation, which are critical in the clearance of pathogens, is scarce in the scientific literature. In this study, we showed a clear role of MALAT1 in macrophage antigen uptake, processing and phagocytosis. A remarkable decrease in the uptake and processing of ovalbumin and phagocytosis of *E. coli* in MALAT1 knockdown macrophages substantiate a close relationship between MALAT1 expression and the vital innate functions of macrophages. Consistent with the pro-M1 function of MALAT1, we observed reduced bacterial clearance in cells silenced for MALAT1. Taken together, our functional assays suggest that MALAT1 knockdown impairs bacterial phagocytosis and clearance, indicating its role in shaping innate immune functions in macrophages.

In addition to initial pathogen/antigen recognition, macrophage activity can be further augmented by secreted cytokines during the course of pathogen clearance. With this intent, we evaluated the secretion of pro- and anti-inflammatory cytokines during the phagocytosis of M1 and M2 macrophages in our study of MALAT1 knockdown. A significant reduction in pro- (IL-1α, IL-1β, IL-6, IL-8, CXCL10, and TNF-α) and anti-inflammatory (IL-1Rα and IL-10) cytokines in MALAT1-ablated M1 and M2 macrophages corroborates attenuated phagocytosis. These results suggest that MALAT1 expression modulates the inflammatory microenvironment by regulating upstream cytokine signaling cascades required for the proper functioning of polarized macrophages. MALAT1 is induced by NFκB activation, and our results show that TLR4/5 ligation upregulates MALAT1 expression in macrophages, suggesting its contribution to positive feedback in the proinflammatory cascade. Indeed, higher MALAT1 expression is associated with inflammation in acute pancreatitis [46]. Wang et al. [47] have also shown that MALAT1 expression induces proinflammatory cytokines by increasing H3 histone acetylation (H3) at the MyD88 promoter. This leads to the induction of IRAK1- and TRAF6-mediated signaling events eventually activating NFκB and downstream proinflammatory cytokines (TNF-α, IL-1β, and IL-6) in microglia causing cerebral injury [47]. Another study demonstrated that upregulation of lncRNA MALAT1 epigenetically represses the expression of anti-inflammatory genes by recruiting PRC2 to their promoters and subsequently induces heightened transcription of inflammatory genes [48]. Overall, MALAT1 expression potentiates a robust innate immune response by promoting cytokine expression through multiple mechanisms and favors the M1 phenotype.

Our findings and previous studies clearly support that MALAT1 controls macrophage polarization; however, the mechanism by which MALAT1 regulates this process remains largely unexplored. We speculate that crosstalk between MALAT1 and miRNA would add another dimension to miRNA-dependent M1/M2 polarization. Several studies have demonstrated the miRNA sponging effect of MALAT1 in biological pathways. Yu et al. [49] showed that MALAT1 could potentially regulate the proliferation and activation of primary hepatic stellate cells by sponging miR-101b and regulating the expression of Rac1 [49]. Another study showed that MALAT1 acts as a competing endogenous RNA (ceRNA) to ZEB2 mRNA and regulates its expression by sponging miR-200 family members [50]. Additionally, MALAT1 was reported to induce the migration and invasion of breast cells by regulating the expression of cdc42 through miR-1 sequestration [51]. In this study, we explored the targets of MALAT1 lncRNAs involved in regulating M1 and M2 polarization. Our lab has previously reported differential expression profiles of miRNAs during macrophage differentiation and function [27, 28]. The miR-30 family includes five members (viz. miR-30a, b, c, d, and e), and all of them are downregulated during monocyte-to-macrophage differentiation [28] and TLR stimulation [38]. The expression pattern of the miR-30 family is antagonistic to MALAT1, suggesting a functional relationship between these noncoding RNAs in macrophage biology. Interestingly, bioinformatics analysis revealed novel binding sites for miR-30b on MALAT1. To date, no report has suggested the role of miR-30b and MALAT1 crosstalk in regulating macrophage polarization. Using dual luciferase assays, we confirmed that miR-30b directly binds to MALAT1. This imperative interaction allows MALAT1 to sequester miR-30b and relieve its cognate targets TLR8 and FcεR1γ from posttranscriptional repression [38; *Naqvi et al., unpublished results*].

miR-30b and MALAT1 are ubiquitous, highly abundant noncoding RNAs. In this study, we show that miR-30b exhibits functional antagonism with MALAT1. Overexpression of miR-30b promotes the M2 phenotype, similar to that observed in MALAT1 knockdown. Our global transcriptome analyses of miR-30b-overexpressing DCs identified hundreds of differentially expressed genes. Several transcription factors, including STAT3, STAT1, PPARG, YY1 and SP1, with well-documented roles in myeloid cell development, monocyte differentiation and M1 and M2 macrophage polarization, were enriched. These results strongly highlight the role of miR-30b in regulating gene networks related to macrophage polarization. In addition to the roles of MALAT1 and miR-30b in orchestrating macrophage polarization events, we evaluated how this noncoding RNA interaction competitively regulates phagocytosis and antigen processing. Compared to MALAT1 knockdown or miR-30b overexpression alone, we observed a remarkably significant reduction in *E. coli* phagocytosis and ovalbumin processing in MALAT1 knockdown along with miR-30b overexpression. These data unequivocally suggest that MALAT1 is an endogenous inhibitor of miR-30b function in macrophages. Functional antagonism between MALAT1 and miR-30 family members is reported to regulate biological processes in other cells. For instance, G. Dong et al. also showed that MALAT1 has the potential to rescue the functional effect of miR-30a *in vitro* and *in vivo* by relieving the inhibitory effects of miR-30a on autophagy in cerebral cortex neurons and middle cerebral artery occlusion-reperfusion (MCAO)-induced ischemic brain infarction in mice, respectively [52]. Another study in primary neurons showed that the MALAT1/miR-30 axis regulates neurite outgrowth in hippocampal neurons by regulating the expression of spastin, a microtubule-severing enzyme important for neurite outgrowth [53]. Taken together, our results show a novel lncRNA:miRNA interaction in macrophages integral to cell plasticity and innate immune responses.

Previous studies have shown that the inflammation caused by periodontal pathogens skew macrophage polarization primarily towards proinflammatory (M1) phenotype, which play a significant role in aggravation of periodontal diseases [40]. Dysregulation of lncRNA and miRNA expression is implicated in various immune-mediated diseases; however the functional relationship between different noncoding RNAs is less explored. Our results show a novel lncRNA and miRNAs interaction that may contribute to the severity of periodontal disease. Given that both MALAT1 and miR-30b are responsive to various inflammatory stimuli, bacterial perturbations in the levels of these highly abundant and ubiquitous noncoding RNAs could play critical role in disease progression. MALAT1 expression promotes the proliferation of human periodontal ligament stem cells [54], as well as regulate their osteogenic differentiation through miR-155-5p [55]. Whereas the knockdown of MALAT1 is reported to inhibit the progression of chronic periodontitis via miR-769-5p [56]. While other studies focused on non-immune cells, our study focused on myeloid macrophages that are central to the pathology of periodontal disease. Importantly, our results show that MALAT1/miR-30b axis regulates the polarization of primary human macrophages and also in inflamed murine gingiva. Higher expression of MALAT1 correlates with increased levels of M1 markers (*iNos2* and *Cox2)* and downregulation of M2 marker *arginase1*. Taken together, our *in vivo* and *in vitro* results support pro-inflammatory role of MALAT1 by promoting M1 phenotype and suppressing M2 phenotype via functional sequestration miR-30b, an anti-inflammatory molecule. Although further studies are required to fully characterize the role of MALAT1/miR-30b axis in periodontal diseases, our findings offer concurrent targeting of different noncoding RNA classes as a novel treatment modality to control periodontal inflammation.

## Conclusion

Our study demonstrates that MALAT1 is responsive to macrophage polarization and activation stimuli and that knockdown of MALAT1 favors the M2 phenotype. Mechanistically, MALAT1 exhibited antagonistic expression and function with another noncoding RNA, miR-30b. The expression of miR-30b is downregulated during macrophage differentiation, and our previous studies have shown that miR-30b is a negative regulator of innate immune responses. In this study, our results suggest that overexpression of miR-30b promotes M2 polarization. By titrating the levels of miR-30b, via a novel binding site, MALAT1 may favor the M1 phenotype. In human and murine healthy and inflamed gingival biopsies we validated antagonistic expression of MALAT1 and miR-30b, which correlates with higher M1 markers in periodontal disease signifying the functional relationship of these noncoding RNA in shaping macrophage phenotype. Concurrent therapeutic targeting of MALAT1 and miR-30b may provide a valuable means of subterfuge to curtail inflammation.

## Acknowledgements

This study was supported by the NIH/NIDCR R01 DE027980 and R03DE027147 to ARN.

## Abbreviations used in this article

APC: antigen-presenting cells
DC: dendritic cells
LncRNAs: long noncoding RNAs
LPS: lipopolysaccharide
Mφ: macrophages
MFI: mean fluorescence intensity
miRNA: microRNA
MPS: mononuclear phagocyte system
mDC: myeloid dendritic cells
PBMCs: peripheral blood mononuclear cells
TLRs: Toll-like receptors
3’UTR: 3’ untranslated region

