## Supplementary Figures for "LncRNA MALAT1/microRNA-30b axis regulate macrophage polarization and function"

**Supplemental Material**

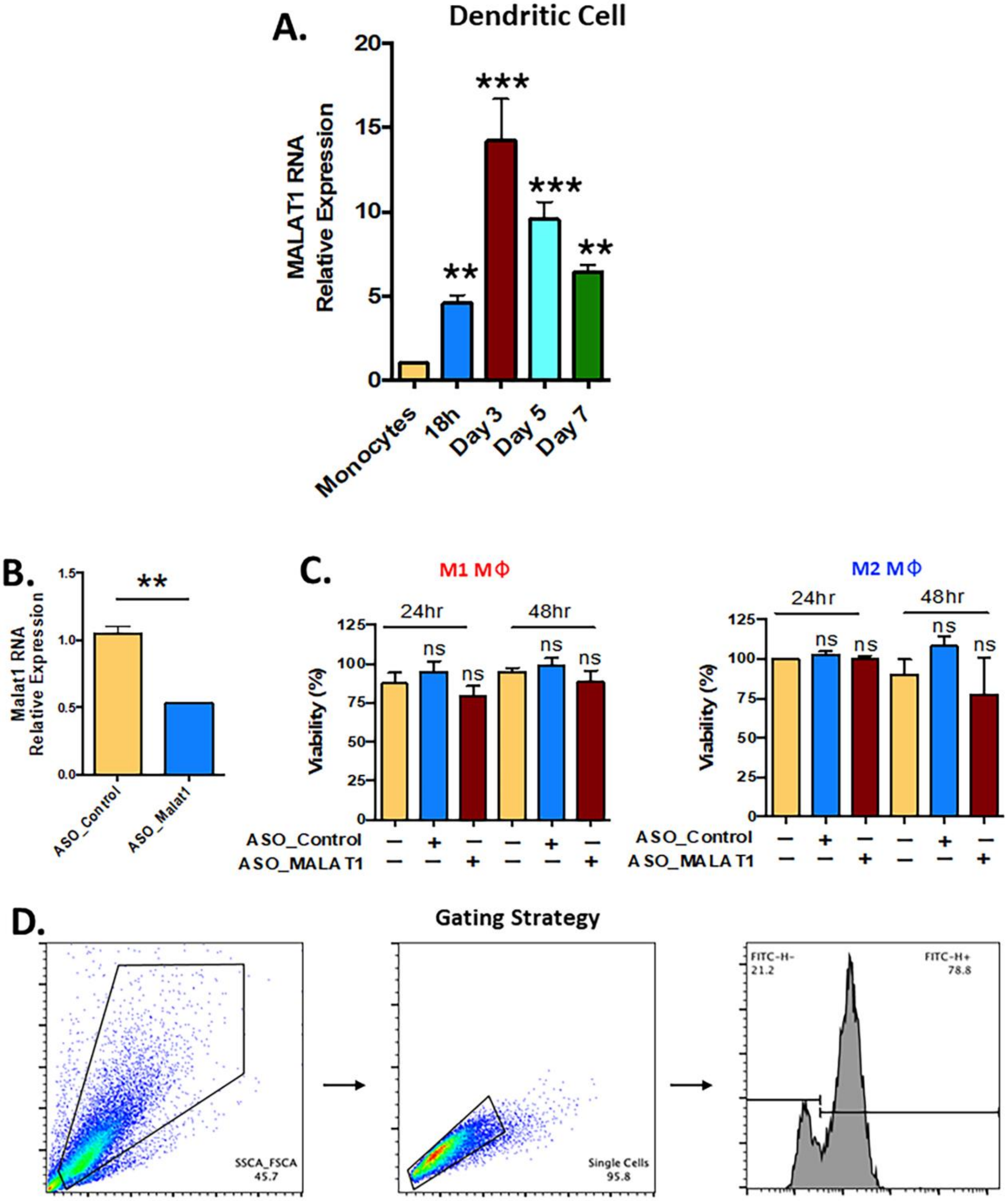


**Figure S1**. **MALAT1 expression, knockdown, impact on cell viability and flow gating**

**strategy.** (**A**) MALAT1 expression is induced during differentiation of monocyte-derived dendritic cells. CD14+ monocytes were differentiated to dendritic cells in the presence of GM-CSF and IL4 (both 50 ng/ml) for seven days. Cells were harvested at indicated time points and RNA was isolated for expression analysis by RT-qPCR. Histogram showing the relative expression of lncRNA MALAT1 in differentiating dendritic cells compared with monocytes. GAPDH was used as an internal control. Data are presented as ±SEM from four donors. **P<0.01, and ***P<0.001 (unpaired, two-tailed Student's t-test). (**B**) Validation of MALAT1 knockdown by GapmeR transfection in M2 macrophages. Cells were transfected with ASO_Control or ASO_MALAT1 (50nM) and total RNA was isolated after 48 h. Expression of MALAT1 was quantified by RT-qPCR. Histograms showing significant reduction in MALAT1 levels compared to control. GAPDH served as endogenous control. Data are presented as ±SEM from four donors in two independent experiments. **P<0.01, and ***P<0.001 (unpaired, two-tailed Student's t-test). (**C**) Cell viability of M1 or M2 macrophages is not impacted by MALAT1 knockdown. Cells were transfected with control or MALAT1 GapmeR (50 nM) and cell viability was assessed by CellTiter 96 AQueous Cell Proliferation Assay at 24 and 48 h post-transfection by recording absorbance (O.D.) at 490nm. Histograms showing percent O.D. calculated compared to Lipofetamine only transfected cells. Data are presented as ±SEM from four donors in two independent experiments. (**D**) Gating strategy used for selecting M1 and M2 macrophages in flow cytometry experiments.

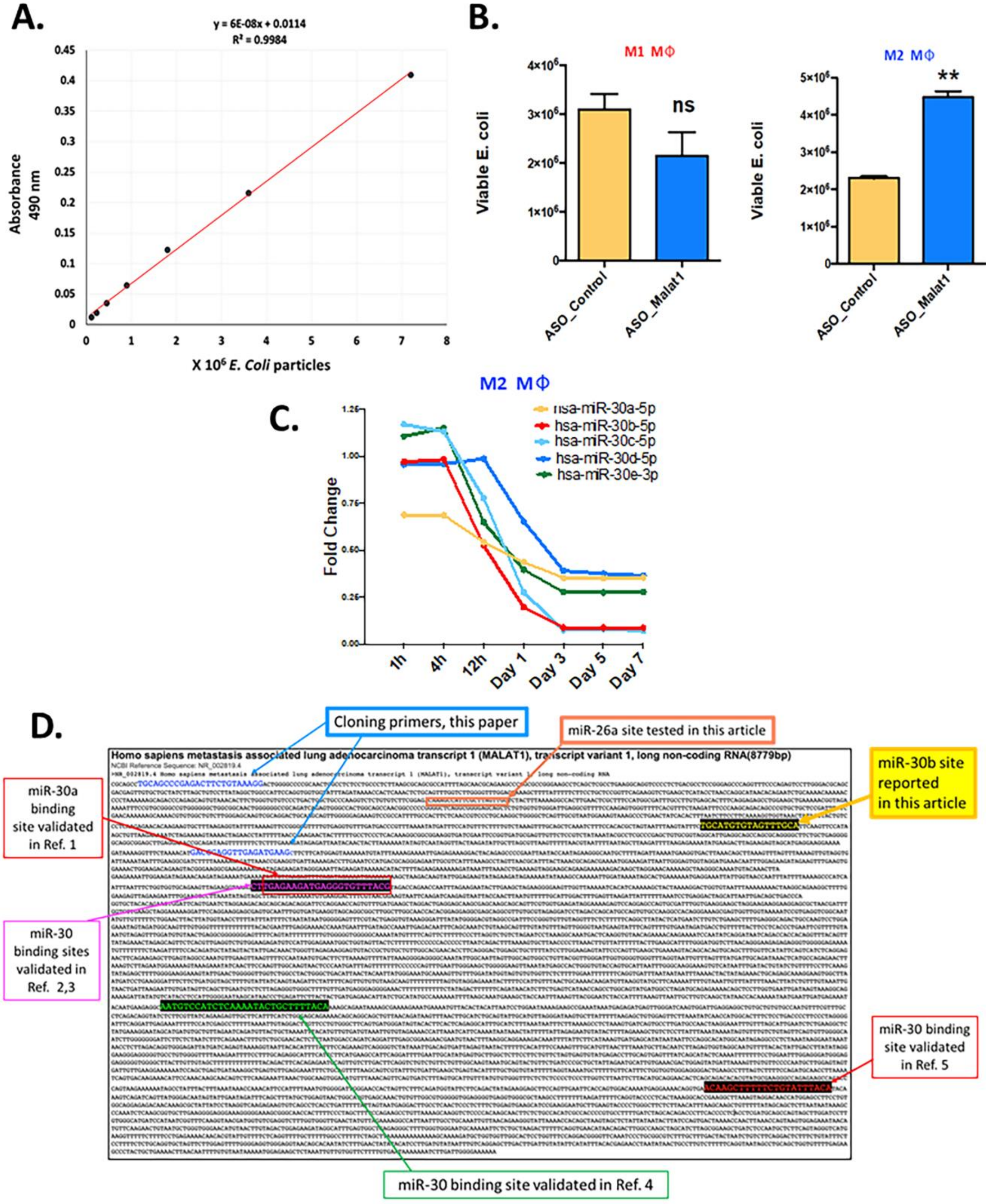


**Figure S2**. **MALAT1 bactericidal assays, miR-30b family expression and novel miR-30b binding sites on MALAT1 RNA.** (**A**) MALAT1 knockdown suppresses bactericidal activity in M2

macrophages. Standard curve of *E. coli* count vs O.D. at 490nm for calculating the unknown *E. coli* particle number in bactericidal activity assay. (**B**) Cells were transfected with control or MALAT1 GapmeRs (50nM) and incubated with live *E. coli* ($5x10^5$ particles resuspended in DMEM) at 10 MOI. After 8 h, cells were lysed and live bacterial quantification was performed as described in methodology. Histogram showing viable *E. coli* numbers in M1 and M2 macrophage in bactericidal activity assay. Data are presented as ±SEM from four donors in two independent experiments. **P<0.01 (unpaired, two-tailed Student's t-test), n.s: non-significant. (**C**) miR-30 family members are downregulated during macrophage differentiation. Expression kinetics of five different miR-30 family members viz., miR-30a-e were plotted from our previously published data microarray data deposited in Gene Expression Omnibus public database (Accession Number GSE60839). (**D**) Schematic highlighting the cloned MALAT1 sequence and novel miR-30b binding sites validated in this study. In addition, we have shown previously validated (see references listed for each site) miR-30 family binding sequence on MALAT1**.**

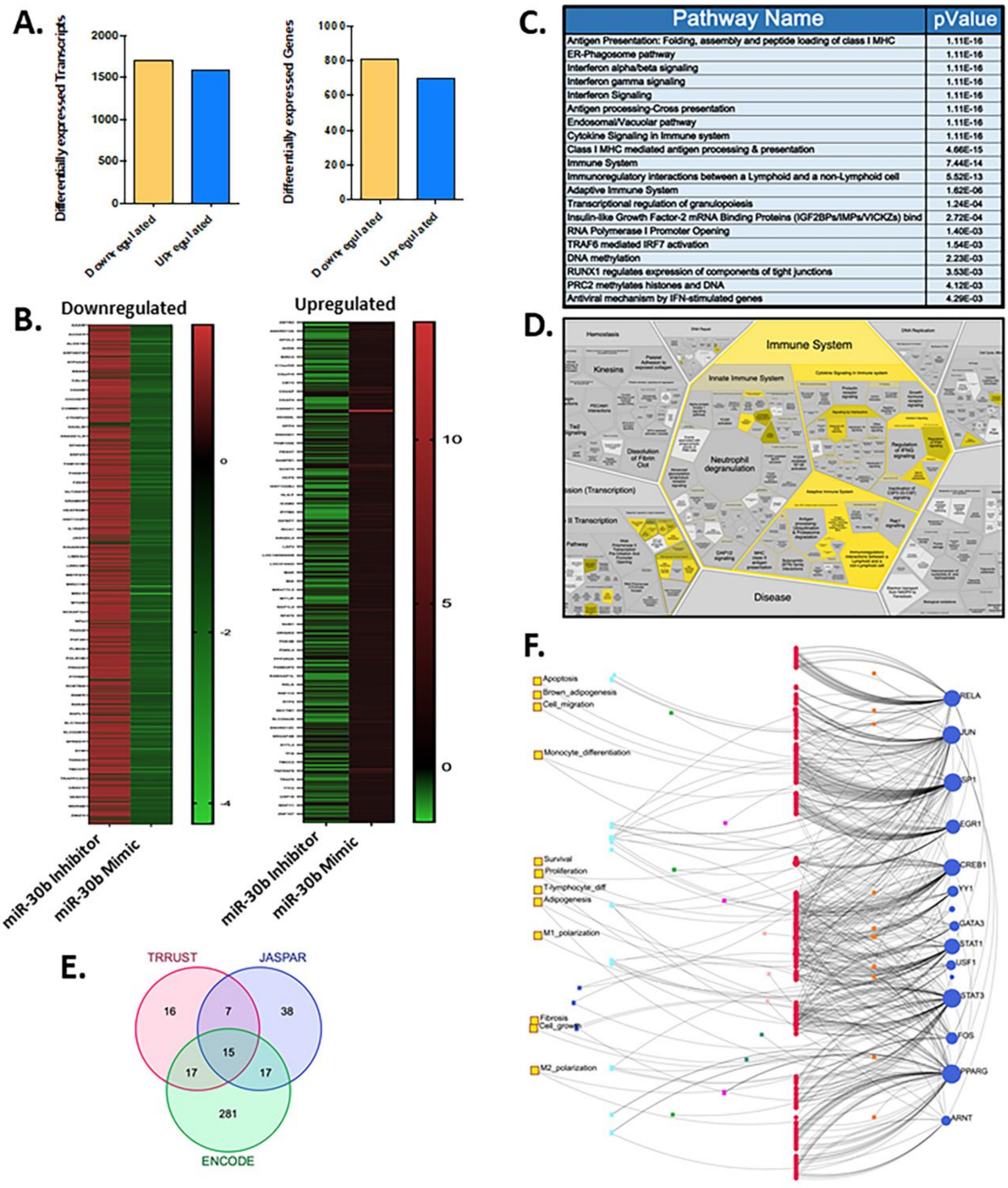


**Figure S3**. **Transcriptome analysis of miR-30b in human CD14+ monocytes-derived dendritic cells (DCs).** Cells were transfected with corresponding control mimic and miR-30p-5p during 36 h. Thereafter, RNA was isolated and transcriptome profiling was performed using microarray analysis (n=3). (**A**) Bar graphs showing the differentially expressed transcripts and genes in miR-30b mimic transfected DCs. (**B**) Heatmaps showing expression profiles of downregulated and upregulated genes in microarray analysis (fold change cut off between −1.25 and 1.25). (**C**) Top 20 enriched pathways identified by reactome analysis of differentially expressed genes. (**D**) Graphical representation of most perturbed pathway in reactome analysis. (**E**) Common transcription factors enriched in different databases. (**F**) Signaling network based on 15 enriched transcription factors.

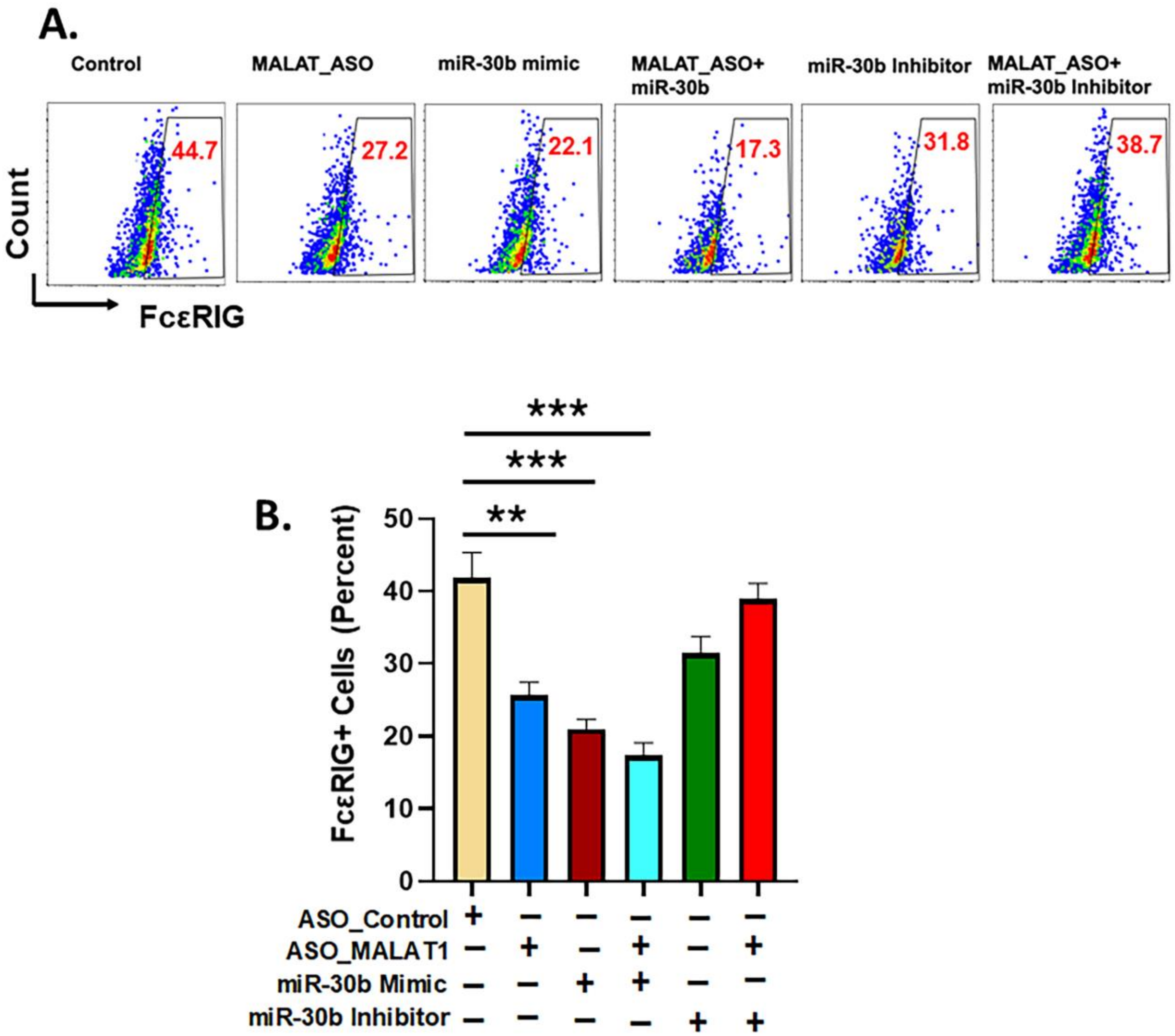


**Figure S4**. **MALAT1 relieves miR-30b target FcεR1G from post-transcriptional suppression.** Cells were transfected with ASO_Control, ASO_MALAT1, miR-30b mimic, miR-30b with ASO_MALAT1, and miR-30b inhibitor with ASO_MALAT1 and the surface expression of FcεRIγ was assessed by flow cytomteric analysis. (**A**) Overlay histograms showing expression pattern of FcεRIγ in different transfections. (**B**) FcεRIγ+ population were plotted as bar graphs. Data are presented as ±SEM from four donors in two independent experiments. *P<0.05 (unpaired, two-tailed Student's t-test), n.s: non-significant.
